# The E3 ubiquitin ligase FBXL6 controls the quality of newly synthesized mitochondrial ribosomal proteins

**DOI:** 10.1101/2022.11.01.514535

**Authors:** Julie Lavie, Claude Lalou, Walid Mahfouf, Jean-William Dupuy, Aurélie Lacaule, Agata Ars, Didier Lacombe, Anne-Marie Duchêne, Anne-Aurélie Raymond, Hamid Reza Rezvani, Richard Patryk Ngondo, Giovanni Bénard

## Abstract

The large majority of mitochondrial proteins is synthesized in the cytosol and then imported to the organelle. To ensure proper mitochondrial functions, the quality of these proteins needs to be guaranteed. Here, we show that the E3 ubiquitin ligase F-box/LRR-repeat protein 6 (FBXL6) participates to the quality of these mitochondrial proteins at the level of the cytosolic translation. We found that lack of FBXL6 has severe effects including mitochondrial ribosomal protein aggregations, altered mitochondrial metabolism and inhibited cell cycle progression in oxidative conditions. FBXL6 was found to interact specifically with ribosomal-associated quality control proteins and chaperones involved in the regulation of newly synthesized proteins and also it preferentially binds newly synthesized mitochondrial ribosomal proteins. Consistently, deletion of the RQC protein, NEMF or HSP70-family chaperone HSPA1A impedes FBXL6 interaction with its substrate. In addition, cells lacking FBXL6 display altered degradation of defective mitochondrial ribosomal protein containing C-terminal alanyl-threonyl extension.

## Introduction

Mitochondria contain about 1200 proteins (Rath et al. 2021). Most of these mitochondrial proteins are encoded by nuclear genes, translated in the cytosol and imported to the various mitochondrial compartments to contribute to the different mitochondrial functions. In mammals, only 13 mitochondrial proteins are directly encoded and translated inside the organelle and they contribute exclusively to the oxidative phosphorylation (OXPHOS) complexes. Translation of nuclear gene-encoded mitochondrial proteins is coupled to their import into the organelle and this is supported by the proximity between the cytosolic ribosomes and the outer mitochondrial membrane (OMM) (Kellems and Butow 1974)(Gold et al. 2017). Lesnik et al. proposed that the cytosolic ribosomes are physically connected to the translocase of the outer mitochondrial membrane (TOM) through specific receptors (Lesnik et al. 2014). Inhibition of the mitochondrial proteins import induces proteostatic response in the cytosol characterized by inhibition of protein synthesis and activation of the proteasomal and protein folding systems (Wrobel et al. 2015)(Boos et al. 2019).

Recently, Izawa et al. have demonstrated in yeast that the VCP/CDC48-associated mitochondrial stress-responsive protein 1 (Vms1) protects mitochondria from toxic aggregations of defective nascent proteins in coordination with the ribosome-associated protein quality control (RQC) pathway. The RQC is a specific mechanism triggered by the ribosome stalling which involves a complex and conserved machinery (Joazeiro 2019). Briefly, Dom34/PELO and Hbs1/HBS1L (yeast/Homo sapiens proteins, respectively) sense the stalled ribosome and recruit Rli1/ABCE1 to disassemble the ribosomal cores (Pisareva et al. 2011). Then, Rqc2/NEMF and Rqc1/TCF25 join the 60S ribosomal subunit and recruit the E3 ubiquitin ligase Listerin (Ltn1/LTN1) to ubiquitinate and target the defective protein for proteasomal degradation (Bengtson and Joazeiro 2010; Defenouillère et al. 2016; Chu et al. 2009). Concomitantly, Vms1 and its mammalian homologue, ANKZF1, are involved in the release of the defective nascent chains from stalled ribosomes (Kuroha et al. 2018)(Verma et al. 2018). This protein was previously reported to be a component of the endoplasmic reticulum- and mitochondrial-associated degradation pathway (ERAD/MAD), playing an important role in the mitochondrial protein stress response in different models (Heo et al. 2010). Thus, the Vms1-dependent mechanism described by Izawa et al. provides a molecular link between the mitochondrial proteostasis and this RQC; this specific process was named, the mitochondrial ribosome-associated protein quality control (mitoRQC) (Izawa et al. 2017). This mitoRQC is essential for mitochondrial functions and yeast strains lacking Vms1 were unable to grow under oxidative conditions showing the importance of this mechanism for mitochondrial metabolic functions (Izawa et al. 2017)(Heo et al. 2010). In mammalian cells, the existence of the mitoRQC has not yet been described but it is highly likely. Paving the way to demonstrating the existence of this process in mammals, Wu et al. showed that mitochondrial damages stall the translation of mitochondrial proteins, leading to the recruitment of PELO and ABCE1 to ribosome/OMM contact sites to promote autophagic mitochondrial degradation (Wu et al. 2018). Furthermore, ANKZF1 was found to be translocated to the mitochondria under oxidative stress (Van Haaften-Visser et al. 2017).

Duttler et al. have proposed that Ltn1 is not the only RQC-associated E3 ubiquitin ligase in yeast and ubiquitination of nascent proteins is carried out by a complex network of E3 ubiquitin ligases (Duttler, Pechmann, and Frydman 2013). Furthermore, this could be the case in higher eukaryotes. Indeed, the levels of E3 ubiquitin ligases in mammalian cells are approximately fourfold to sixfold higher than those of yeast cells suggesting a higher specificity regarding substrate recognition or/and physiological regulation (Li et al. 2008)(Finley et al. 2012). Recently, it was found that NEMF can target RQC substrates for ubiquitin-dependent degradation independently of Listerin and that this degradation involves the E3s, CRL2^KLHDC10^ and Pirh2 (Thrun et al. 2021). In this line, our current study investigated the role of E3 ubiquitin ligases belonging to the F-box/Leucine rich-repeat family (FBXL). This family includes FBXL4, which is responsible for a mitochondrial syndrome characterized by decrease expression of mitochondrial proteins and mtDNA depletion and impaired mitochondrial metabolic functions (Gai et al. 2013)(Bonnen et al. 2013)(Huemer et al. 2015). In this study, we showed that, like FBXL4, the FBXL6 loss of function is responsible for impairment of the mitochondrial metabolism. Yet, FBXL4 displays in mitochondrial localization while FBXL6 is located to the cytosol and to the nucleus. We found that both E3 ubiquitin ligases interact with RQC proteins and chaperones associated with mitochondrial proteostasis and that FBXL6 specifically interacts with mitochondrial ribosomal proteins. Accordingly, we propose that FBXL6 participates to the quality control of neosynthesized mitochondrial proteins in coordination with cytosolic ribosome quality control hub.

## Results

### FBXL4 and FBXL6 are implicated in mitochondrial biogenesis during the cell cycle

FBXL4 is a E3 ubiquitin ligase that mutations are responsible for mitochondrial disease characterized by inhibited mitochondrial energy metabolism (Gai et al. 2013)(Bonnen et al. 2013)(Huemer et al. 2015). Consistently, our results revealed that CRISPR/Cas9-mediated deletion of this enzyme in Hela cells (FBXL4-KO) specifically impaired the mitochondrial metabolism since FBXL4-KO cell growth was inhibited in oxidative conditions but not in glycolytic conditions (Fig. 1A-B, S1A-B) (Reitzer, Wice, and Kennell 1979)(Melser et al. 2013). Interestingly, we found that cells carrying deletion of FBXL6, another FBXL family, displayed the same cell growth discrepancy suggesting a mitochondrial metabolic dysfunction (Fig. 1A-B, Fig. S1C-D). This effect was compensated by FBXL6 ectopic expression in KO cells (Fig. S1E). Confirming these results, we found that both FBXL4- and FBXL6-KO cells were arrested at the G2/M transition under oxidative conditions but not when cultured in medium containing glucose (Fig. 1C-D). Furthermore, we also measured impact of these deletions on mitochondrial bioenergetics by measuring the oxygen consumption rates (OCR) and mitochondrial ATP production (Fig. 1E-F). Both FBXL4- and FBXL6-KO cells displayed moderate decrease of endogenous OCR, ranging from 8.5 to 22.6% of the control OCR (Fig. 1E, S1F), whereas mitochondrial ATP production was inhibited by 25 to 40% in FBXL4-KO cells and by about 30% in FBXL6-KO cells (Fig. 1F). These results support a decrease in the coupling efficiency between oxygen consumption and ATP production. Since implication of FBXL6 regarding mitochondrial functions, we investigated whether FBXL6 deletion induced changes in biological pathway by comparing the whole proteome of FBXL6-KO cells and control cells (Fig. 1G, Table S1). We reported increased expression of proteins involved in the ribosome biogenesis pathway (Fig. 1H, Table S1). Concomitantly, using immunoblots, we found that FBXL6 deletion was associated with decreased levels of cytochrome c oxidase subunit 2 (mtCO2) and, notably, higher levels of some mitochondrial proteins, such as NDUFS3 or the mitochondrial ribosomal protein L45 (MRPL45) (Fig. S1G-H).

**Figure 1.**
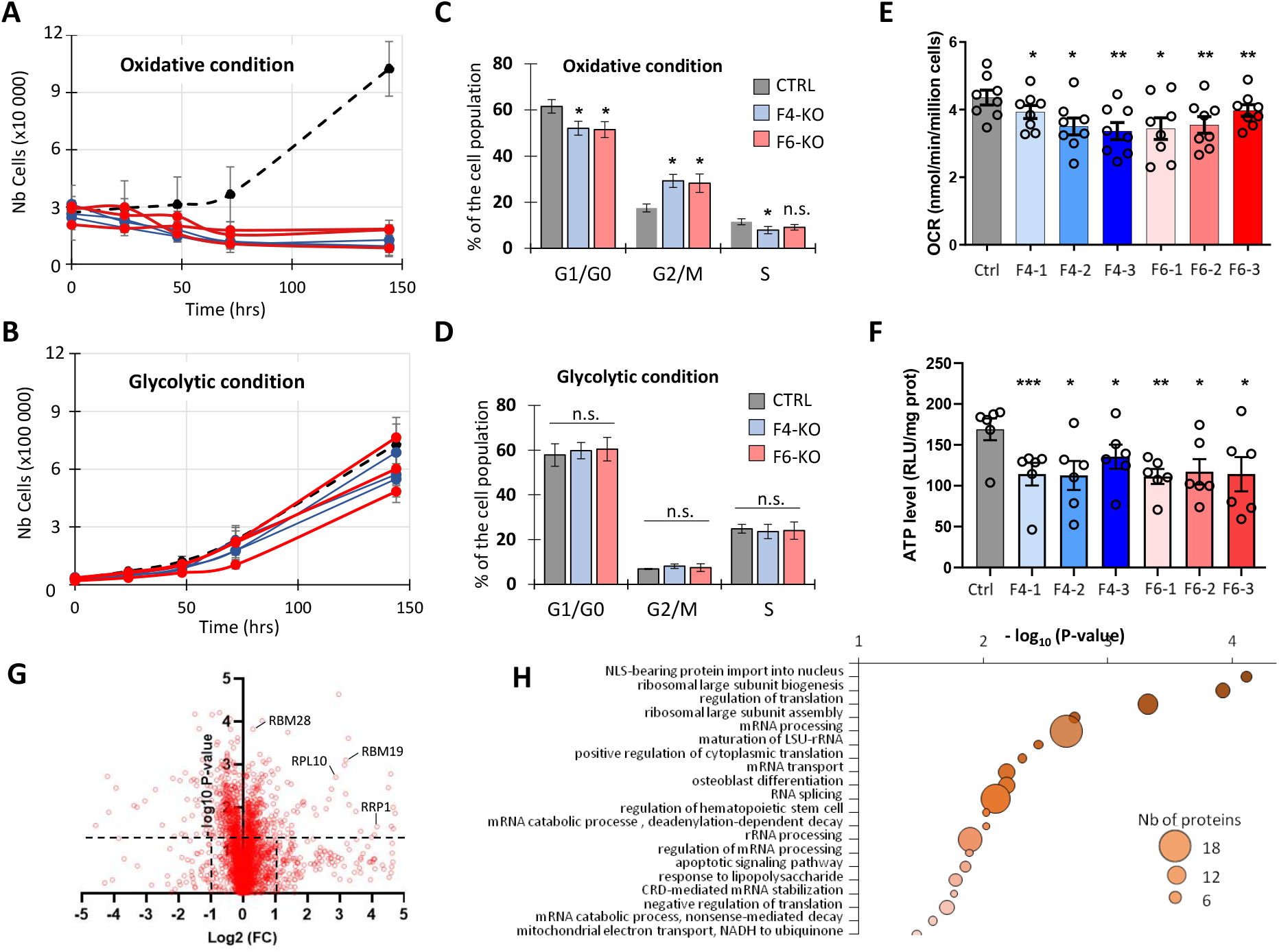
Knockout of F-box/LRR-repeat protein 4 (FBXL4) and FBXL6 causes mitochondrial defects and cell cycle perturbations. (A-B) Growth of FBXL6-knockout (KO) (red), FBXL4-KO (blue), and control (black) cells were measured under glycolytic (A) and oxidative (B) conditions. Growth was measured in three different FBXL4- or FBXL6-KO Hela cell clones (means ± SEM, n = 3). (C-D) Cell cycle analyses performed on FBXL4-KO, FBXL6-KO, and control cells grown under oxidative (C) or glycolytic (D) conditions (means ± SEM, n = 4-8, * P<0.05, ANOVA Kruskal-Wallis test). (E-F) Basal oxygen consumption rates (OCR) (E) and mitochondrial ATP levels (F) of FBXL4-KO (blue shade), FBXL6-KO (red shade) and control (black) cells. Bars indicate the means ± SEM for the different KO clones and open circles represent single measurements. * P<0.05, ** P<0.01, RM ANOVA-Holm-Sidak’s multiple comparisons test. For OCR, n = 8 and for ATP level, n = 6. (G) Proteomic analyses of FBXL6-KO and control whole cell extracts cells by using mass spectrometry. Graphs represent the volcano plots obtained by comparing the differential protein expression between FBXL6-KO and control cells (G). Each symbol represents an identified protein. (N = 3). Examples of proteins involved in the mitochondrial functions or in the ribosome biogenesis are cited. (H) Gene ontology enrichment analyses of up regulated proteins in FBXL6-KO vs. control cells. Pathways are represented according to −log10 of p-value (abscise axis). The circle size is proportional to the number of proteins.

### FBXL6 is not an intrinsic mitochondrial E3 ubiquitin ligase

Since it was reported that FBXL4 is a mitochondrial E3 ligase (Gai et al. 2013)(Bonnen et al. 2013), we analyzed the cellular distribution of FBXL6 using immunofluorescence (Fig. 2A). Expectedly, we found that myc-tagged FBXL4 localized to the mitochondria (Pearson’s coefficient Myc vs. TOMM20 = 0.90±0.01) (Fig. 2B). In contrast, FBXL6 displayed a diffused localization in the cytosol and in the nucleus (Fig. 2A-B, Pearson’s coefficient Myc vs TOMM20 = −0.01±0.02). The nuclear localization represented 46,8 % ± 3,5 of the cell population and 53,21%±3,6 for the cytosolic distribution (N=5, P-value = 0.24, unpaired t-Test). In comparison, FBXL3, another E3 ubiquitin ligase from the same family displayed a strictly nuclear localization (Fig. 2A-B), as previously reported (Hirano et al. 2013). Interestingly, in silico analyses of FBXL4 protein sequences revealed a putative mitochondrial leading sequence (MLS) at the N-terminal of FBXL4 and surprisingly, these analyses also predicted that FBXL6 could be a mitochondrial protein (Fig. S2). Thus, we assayed the presence of a leading sequence by coupling the N-terminal amino acids of FBXL4 (1-25aa) and FBXL6 (1-20aa) to green fluorescent protein (GFP). We demonstrated that the FBXL4 sequence targeted the GFP to the mitochondria while the FBLX6 sequence failed to address the fluorescent protein to any specific subcellular compartment (Fig. 2C-D). Using cell fractionation assays, we confirmed that FBXL6 had a spread subcellular localization (Fig 2E). Nevertheless, a part of FBXL6 was retained to the mitochondrial purified fraction free of associated membranes (Fig. 2E). This part of FBXL6 was quickly digested during a trypsin accessibility assay, suggesting that it was actually loosely associated to the organelle surface, facing the cytosol (Fig. 2F).

**Figure 2.**
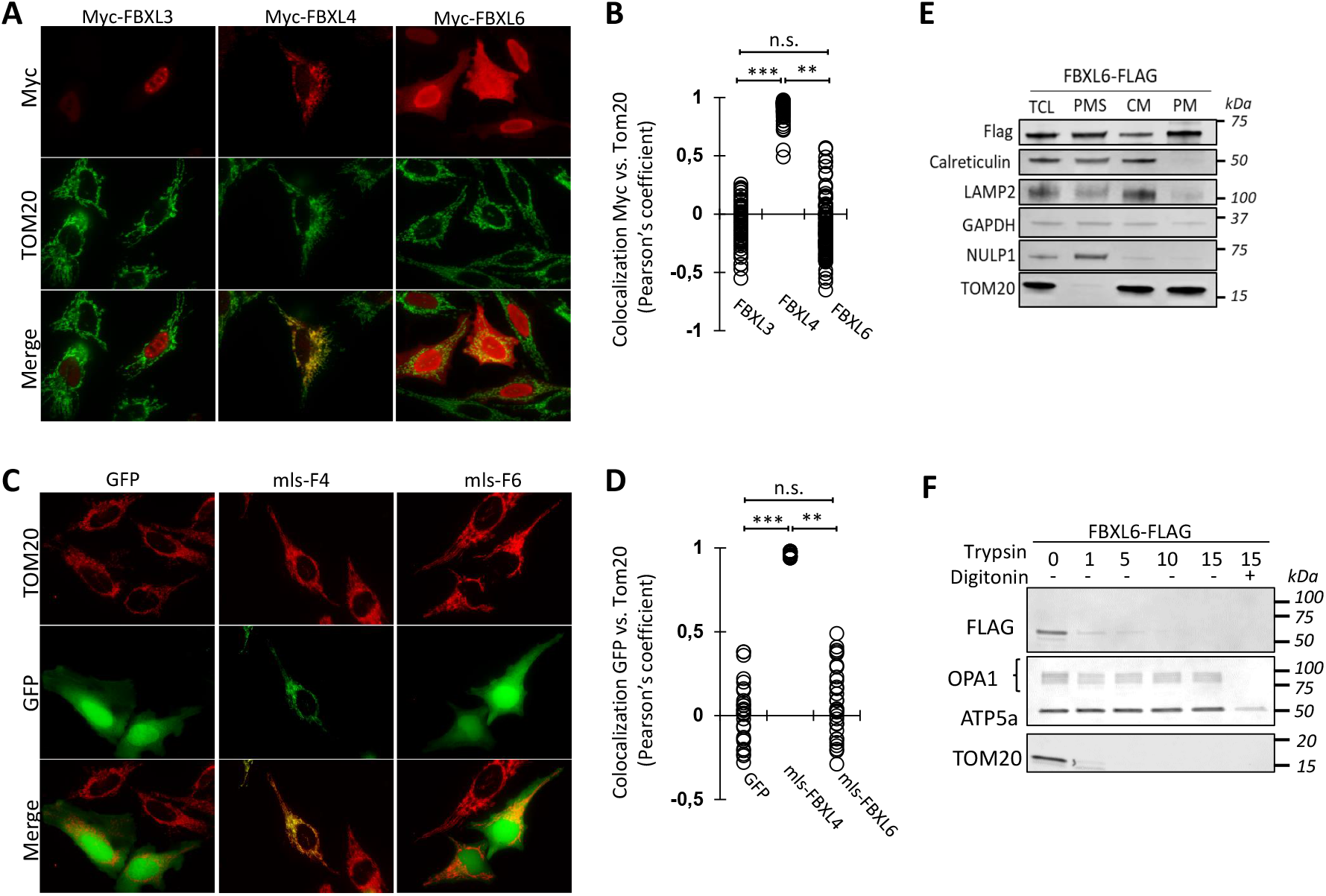
F-box/LRR-repeat protein 6 (FBXL6) is located to the cytosol and the nucleus. (A) Immunofluorescence performed on Hela cells ectopically expressing myc-tagged FBXL3, FBXL4 or FBXL6. FBXL mitochondria were labeled using anti-myc (red) and anti-translocase of the outer mitochondrial membrane 20 (TOMM20) (green) antibodies, respectively. (B) Quantification of FBXL localization to mitochondria obtained from (A). Pearson’s correlation coefficient between the TOMM20 and myc staining; each dot represents the value for one cell (N = 4-6, 70-100 cells, *** P < 0.001 ANOVA, Kruskal-Wallis test) (C) Immunofluorescence performed on Hela cells expressing green fluorescent protein (GFP) fused to the theoretical mitochondrial leading sequence of FBXL4 or FBXL6. Mitochondria were labelled using an anti-TOMM20 antibody. (D) Quantification of the chimeric GFP targeted to the mitochondria from (C). Each dot represents the Pearson’s correlation coefficient for single cells. (N = 3, 40-60 cells, *** P < 0.001 ANOVA, Kruskal-Wallis test) (E) Immunoblots performed using the total cell lysate (TCL), Post mitochondrial supernatant (PMS), crude mitochondria (CM) and purified mitochondria (PM). Fractions were obtained from HEK cells expressing FBXL6-Flag. FBXL6 was detected using anti-Flag. Calreticulin, LAMP2, GAPDH, NULP1 and TOM20 were used as marker of the ER, lysosomes, cytosol, nucleus and mitochondria, respectively. (F) Immunoblots showing the results of the trypsin accessibility assay performed on mitochondria isolated from HEK cells expressing FBXL6-Flag. OPA1 mitochondrial dynamin like GTPase (OPA1), ATP5A, and TOMM20 were used as respective markers of the intermembrane space, and the inner and the outer mitochondrial membranes.

### FBXL6 regulates newly synthesized ribosomal proteins

To understand FBXL6-associated mechanism, we analyzed its interactome by using coimmunoprecipitation coupled to mass spectrometry. FBXL6 interacted with 451 proteins including Skp1, Rbx1 and Cullin1 (Fig. 3A, Table S2) confirming that FBXL6 can form the SCF complex (SKP1-Cullin1-FBXL6) like other FBXLs. FBXL6 interacting proteins included also numerous chaperones and co-chaperones involved in the folding and transport of newly synthetized proteins (e.g. HSPA8, HSPA1A/B, HSP90AB1, BAG2) and involved in delivering of neosynthesized protein to mitochondria (HSP90AA1) (Young, Hoogenraad, and Hartl 2005). Of note, we found interactions with the RQC proteins, NEMF and PELO (Fig. 3A)(Pisareva et al. 2011)(Shao et al. 2015). In addition, we also noted interactions with cytosolic and mitochondrial ribosomal proteins. Overall pathway analyses supported that FBXL6 contributes to the protein folding and to the ribosomal protein biogenesis including mitochondrial ones (Fig. 3B, TableS3).

**Figure 3.**
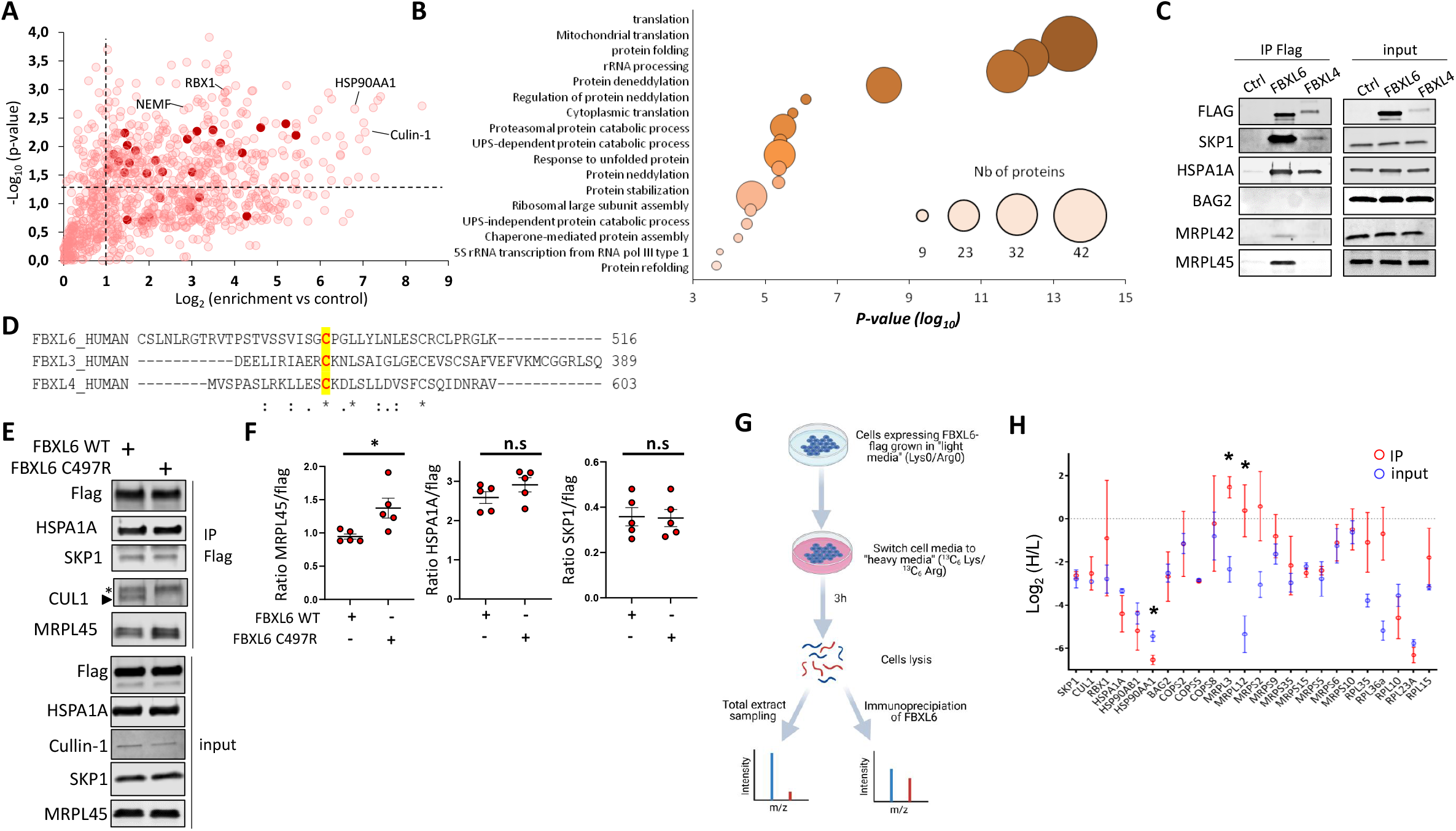
Identification of FBXL6-interacting proteins. (A) HEK cells were transfected with β-gal (ctrl) or FBXL6-Flag. After Flag-immunoprecipitation (IP), interacting proteins were identified using mass spectrometry. Plots representing the fold changes (log2) and p-values (-log10) for proteins interacting with FBXL6 vs. control. The names of proteins of interest are specified. Red dots are mitochondrial ribosomal proteins. N= 3 independent IPs. (B) Gene ontology enrichment analyses performed on FBXL6 interacting proteins. Enriched pathways are displayed with log10 of p-value (abscise axis). The circle size is proportional to the number of counted proteins. (C) HEK cells transfected with β-gal (control), FBXL4-Flag, and FBXL6-Flag and IP performed using an anti-Flag antibody. Immunoprecipitated proteins were analyzed using immunoblots (N = 6). Interactions with proteins were analyzed by immunoblots (see details in the panel). D) FBXL3, 4 and 6 amino acids sequences of alignment. Substrates binding domains represented and a conserved cysteine highlighted. (E-F) Immunoprecipitation of WT- or C497R-FBXL6 flag. WT- or C497R-FBXL6 flag were expressed in HEK cells and interactions with various SKP1, HSPA1A and MRPL45 were analyzed by immunoblots. Graphs represent quantification immunoblots and values are normalized to the Flag level (F). (N =5, * P<0.05, unpaired t-test) (G) Pulsed SILAC experimental design. HeLa cells expressing FBXL6 grown in light (L, R0K0) media were transferred to heavy (H, R10K8) media for 3h. Samples of the total fractions (input) were collected and immunoprecipitations of FBXL6 (IP) performed in parallel. The H/L ratios were determined by mass spectrometry. (H) H/L ratio determined for FBXL6 interacting proteins in input (Blue) or after immunoprecipitation (Red). Ratio were expressed as log2. Blue and red bars are corresponding to H/L ratio measured in input or IP fractions, respectively. (N = 3, * P < 0.05, unpaired T-test).

Implications of FBXL6 in these pathways were further validated using others approaches. First, by immunoblots coupled to immunoprecipation, we showed that FBXL6 and also FBXL4 interacted with SKP1 or HSPA1A. By this method, we also confirmed that FBXL6 binds mitochondrial ribosomal proteins such MRPL45 and MRPL42 and it was not the case for other members of the FBXL family like FBXL4 or FBXL3 (Fig. 3C, S3A). Secondly, because E3 ubiquitin ligases have labile and transient interactions with their substrates and partners, we analyzed FBXL6 interactome using a biotin proximity labelling assay based on TurboID (Branon et al. 2018). HCT116 cells expressing stably GFP-TurboID or FBXL6-TurboID (Fig. S3B-C) were treated with biotin for 0.5 or 16 h (Fig. S3D-F). Biotinylated proteins were identified by mass spectrometry after streptavidin pull down (Fig S3E-F). Pathways analyses also confirmed that FBXL6 interacts with mitochondrial ribosomal proteins and proteins involved in the folding process (Fig. S3G, table S4).

We further analyzed FBXL6 interacting properties by targeting its substrate binding domain. Indeed, FBXL6, 3 and 4 contains a conserved cysteine in the substrate binding domain (Fig 3D)(Mason and Laman 2020) and mutations of this cysteine to an arginine in FBXL4 (C584R) and FBXL3 (C358R) are responsible for specific rare diseases (Sabouny et al. 2019)(Ansar et al. 2019). We found that mutation of this conserved amino acid in FBXL6 (C497R) did not affect FBXL6 binding to HSPA1 or SKP1 (Fig 3E-F). However, it increased binding of the substrate MRPL45 (Fig 3E-F). These results suggested that the C497R mutation inhibits the ubiquitin transfer from E2 to the MRPL45 hampering the substrate release from the E3. This obstruction induced the release of SKP1-FBXL6 from the SCF since FBXL6 (C497R) didn’t associate with Cullin-1 (Fig 3E-F).

As we showed above FBXL6 is not a mitochondrial E3 (see Fig.2) and accordingly, we postulated that FBXL6 interactions with mitochondrial ribosomal proteins occurred in the cytosol with neosynthesized proteins. To assess this hypothesis, we analyzed whether FBXL6 had preferential interactions with neosynthesized proteins using a stable isotope labeling by amino acids in cell culture approaches (SILAC). Cells expressing FBXL6 were incubated for 3h with heavy SILAC media and we measured enrichment of labeled proteins in both input and FBXL6 immunoprecipitated fractions (Fig 3G). After SILAC treatment, proteins have incorporated about 10% of heavy amino acids and this level was not affected either by IP procedures or by the FBXL6 expression (Fig. S3H). When we compared labeling levels of FBXL6 constitutive partners like SKP1, Cullin1 or HSPs proteins, we measured no difference before and after immunoprecipitation (fig 3H). In contrast, several cytosolic or mitochondrial ribosomal proteins such RPL35, RPL36a, MRPL3 and MRPL12 found in IP fractions were enriched in heavy amino acids labeling as compared to inputs (Fig. 3H). These results supported that FBXL6 interacted preferentially newly synthesized proteins.

### FBXL6 is involved in the Ribosome associated Quality Control

Identification of FBXL6 interactome as well as its preferential binding to newly synthesized proteins suggest that this E3 participates to protein quality control associated to the translation. To test this hypothesis, we generated a GFP carrying an (Ala-Thr)10 extension at the C-terminus to mimic CAT-tailed proteins, which are obtained upon activation of the RQC (Shen et al. 2015)(Kostova et al. 2017). These chimeric proteins were targeted either to the mitochondria (mtGFP-CAT) or to the cytosol (cGFP-CAT). Like previously reported by Izawa et al. in yeast model (Izawa et al. 2017), cGFP-CAT formed SDS-resistant aggregates and notably, this was also the case of mt-GFP-CAT (Fig S4A). Inhibition of the proteasome promoted these aggregates (Fig S4A). Regarding the cellular localization, cGFP-CAT aggregated in precise foci within the cytosol whereas the control cGFP displayed a homogenous cytosolic distribution (Fig. S4B). We observed that mtGFP-CAT also formed aggregates and that these aggregates were located to the mitochondria (Fig. S4C). Epoxomicin-induced proteasome inhibition dramatically increased the formation of aggregates in the cytosol but also in the mitochondria. Accordingly, the RQC and the ubiquitin-dependent degradation also prevent accumulation defective newly synthesized mitochondrial proteins in mammalian cells.

Because GFP are xenoprotein that are particularly stable in mammalian cells, we developed similar approach using the FBXL6 endogenous substrate, MRPL45. We have generated a CAT-tailed and a stop codon deleted MRPL45 (MRPL45-CAT or –NS, respectively). First of all, these defective MRPL45 were highly unstable as compared to WT MRPL45 or to GFP-based constructs (Fig. 4A). Then, we found that expression of MRPL45-CAT and MRPL45-NS induced translocation of FBXL6 to precise cytosolic foci and the degree of colocalization between the E3 and defective proteins significantly improved (Fig. 4BC). In contrast, FBXL6 displayed a spread cytosolic distribution and no colocalization with the mitochondrial ribosomal protein was observed in cells expressing WT-MRPL45 (Fig. 4B-C). These results suggested that presence of altered ribosomal proteins mimicking a defective translation activated the FBXL6-dependent mechanism.

**Figure 4.**
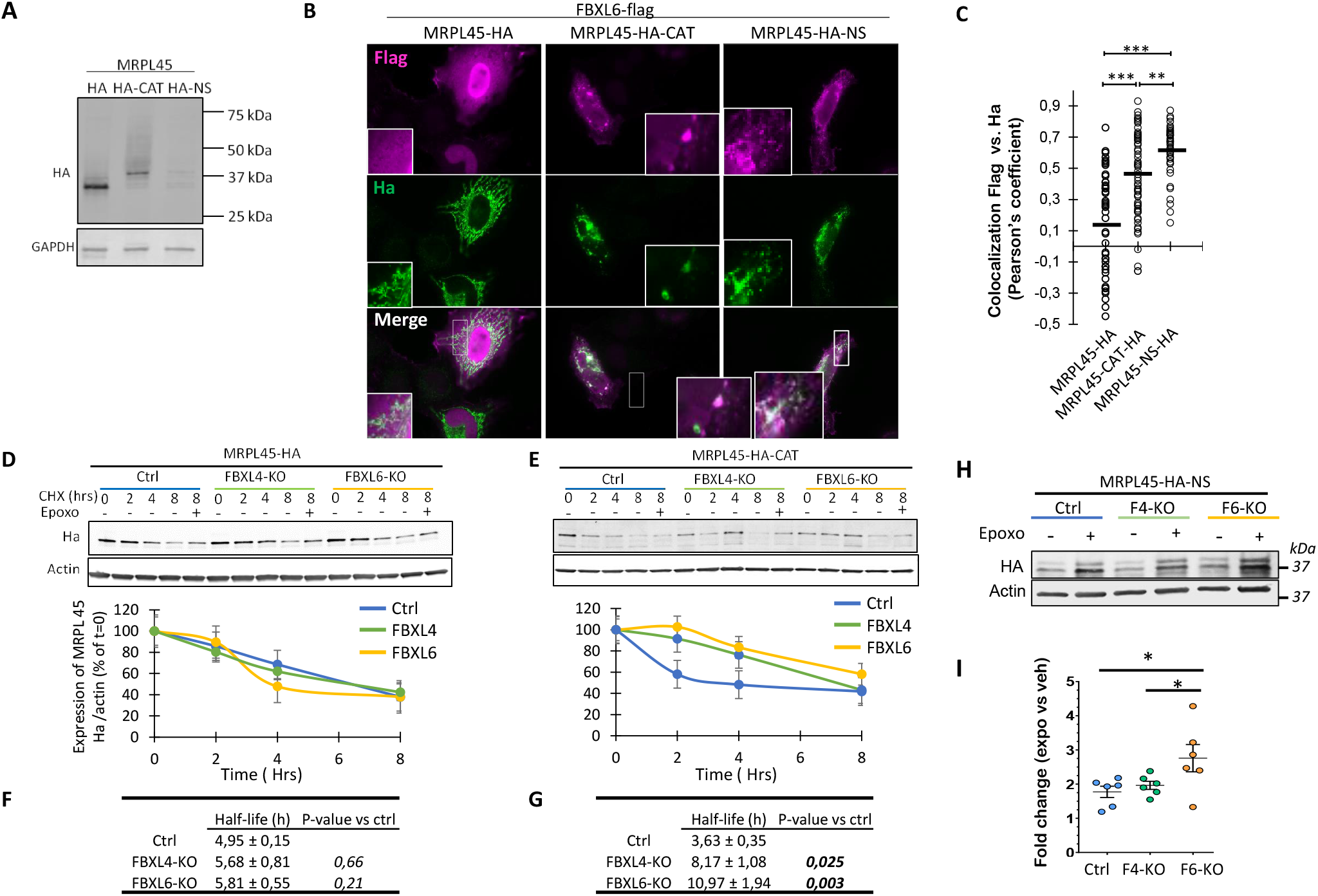
F-box/LRR-repeat protein 6 (FBXL6) is involved in the degradation of defective MRPL45. (A) Immunoblot showing expression of MRPL45-HA, MRPL45-HA-CAT and MRPL45-HA-NS in HEK cells using anti-Ha antibody. (B) Immunofluorescence performed on Hela cells expressing FBXL6-Flag and transfected with mitochondrial ribosomal protein L45 (MRPL45)-hemagglutinin (HA), MRPL45-HA-C-terminal alanine and threonine (CAT), or MRPL45-HA-lacking a stop codon (NS). MRPL45 and FBXL6 were labelled using anti-HA and anti-Flag antibodies, respectively. (C) MRPL45 and FBXL6 colocalization was quantified by measuring the Pearson’s correlation coefficient between HA and flag labelling. Each dot is the value for one cell (N = 3-4, 60-120 cells, *** P < 0.001, ** P < 0.01, ordinary 1W ANOVA, Holm-Sidak test). (D-E) FBXL6-KO, FBXL4-KO, and control cells were transfected with MRPL45-HA (D) or MRPL45-HA-CAT (E) and treated with cycloheximide for 0h, 2h 4h and 8h. The protein levels of MRPL45-HA and MRPL45-HA-CAT were evaluated using immunoblots. The plots show protein quantification normalized to that of t = 0 (N = 4). (F-G) The calculated half-life of each protein is summarized in the table under the graph. First order approximation was used for this calculation. (N = 4, RM ANOVA, Fisher test). (H-I) Accumulation of MRPL45-HA-NS was analyzed in FBXL6 KO, FBXL4-KO and control cells. Cells expressing MRPL45-HA-NS were treated with epoxomicin for 8 h. Level MRPL45-HA-NS were revealed by immunoblots using anti-Ha (H). MRPL45-HA-NS level were normalized to actin and expressed as fold change of untreated control cells. (N=6, * P-value <0.05, Two Way-Anova)

Next, we tested whether FBXL6 is involved in the degradation of these proteins. Degradation rate of WT-MRPL45 was not significantly impacted by deletion of FBXL6 (Fig. 4D). MRPL45-CAT was promptly eliminated in a proteasome-dependent manner in control cell while lack of FBXL6 hampered the degradation of this protein (Fig. 4E). Accordingly, we measured that half-life of MRPL45-CAT increased by 3-fold in FBXL6-KO as compared to control cells (Fig. 4F-G). We also measured inhibition of degradation in FBXL4-KO cells (Fig. 4D-G). Similar data were obtained when we compared degradation rate of MRPL42-WT vs MRPL42-CAT in WT and FBXL6-KO cells (Fig. S5A-B). To demonstrate implications of FBXL6 in the specific degradation of defective mitochondrial ribosomal proteins, we performed same experiment using a CAT-tailed version of the subunit A of the mitochondrial succinate dehydrogenase (SDHA-CAT) and no difference of degradation were found between control, FBXL4-KO or FBXL6-KO cells, (Fig. S5C). Regarding MRPL45-NS, the defective protein was very rapidly degraded and displayed a very short half-life making it difficult to determine the effect of the deletion of FBXL6 (Fig. S5D). However, we found that deletion of FBXL6 potentiated the accumulation of MRPL45-NS induced by proteasome inhibition, supporting that FBXL6 participated also to the proteasome-dependent degradation of the non-stop protein (Fig. 4H-I).

### Implication of RQC proteins and chaperones in the FBXL6-dependent mechanism

To better understand FBXL6-dependent mechanism, we analyzed the role played by FBXL6 partners. First, co-immunoprecipitation coupled to immunoblots confirmed that FBXL6 interacts specifically with proteins involved in the initial steps of the RQC including ZNF598, PELO and in lesser extent NEMF (Fig. 5A). In contrast, no interactions were observed with proteins associated to the degradation of defective nascent proteins such as LTN1, ANKZF1, TCF25 or VCP (not shown). Accordingly, we analyzed if inhibition of these initial and degradation steps promoted FBXL6-dependent mechanism. Interesting, we found that knockdown of NEMF decreased the endogenous level of MRPL45 in FBXL6-KO vs. control cells suggesting FBXL6 and NEMF had coordinated actions (Fig. S6A). No significant effects were observed upon deletion of knockdown LTN1. Then, we analyzed the role of these RQC proteins regarding FBXL6 ability to bind MRPL45. When the sensing of ribosome obstruction is abolished by NEMF deletion, we found that FBXL6 ectopic expression is reduced. Besides, interactions between FBXL6 its substrate, MRPL45, or its partner, HSPA1A were specifically inhibited in NEMF-KO but it was not fully abolished (Fig 5B-C). Data obtained through transient silencing shown that other RQC proteins are susceptible affected interaction with MRPL45 (Fig S6B). Regarding the degradation step, deletion of LTN1 didn’t affect interactions of FBXL6 with either MRPL45 or HSPA1A but it strengthened association with SKP1 (Fig 5B-C). Using LTN1/NEMF double KO, we demonstrated that this effect was prevented by lack of NEMF. Next, we analyzed FBXL6 localization upon deletion of NEMF, LTN1 and NEMF/LTN1. Deletion of these proteins modified FBXL6 subcellular localization by inducing its relocation to cytosolic cluster like previously observed upon expression of mrpl45-CAT and MRPL45-NS (Fig 5D-E, see also fig. 4B). Notably, we didn’t observed relocalization of GFP when expressed in these KO cells supporting that this clustering dependent (Fig 5E, S6C). To analyze if FBXL6 and RQC proteins act synergistically, control and NEMF-KO cells expressing MRPL45-NS were treated with epoxomicin. We found that MRPL45-NS accumulated over the time in control cells (Fig. 5F). Notably, in NEMF-KO cells, we observed that this accumulation was strongly reduced over the time and included build up of lower molecular form of NS-MRPL45 like previously reported (Shen et al. 2015). This effect was reversed by expression of FBXL6 showing that the FBXL6-dependent mechanism can bypass the sensing of the 60S obstruction by NEMF (Fig. 5F).

**Figure 5.**
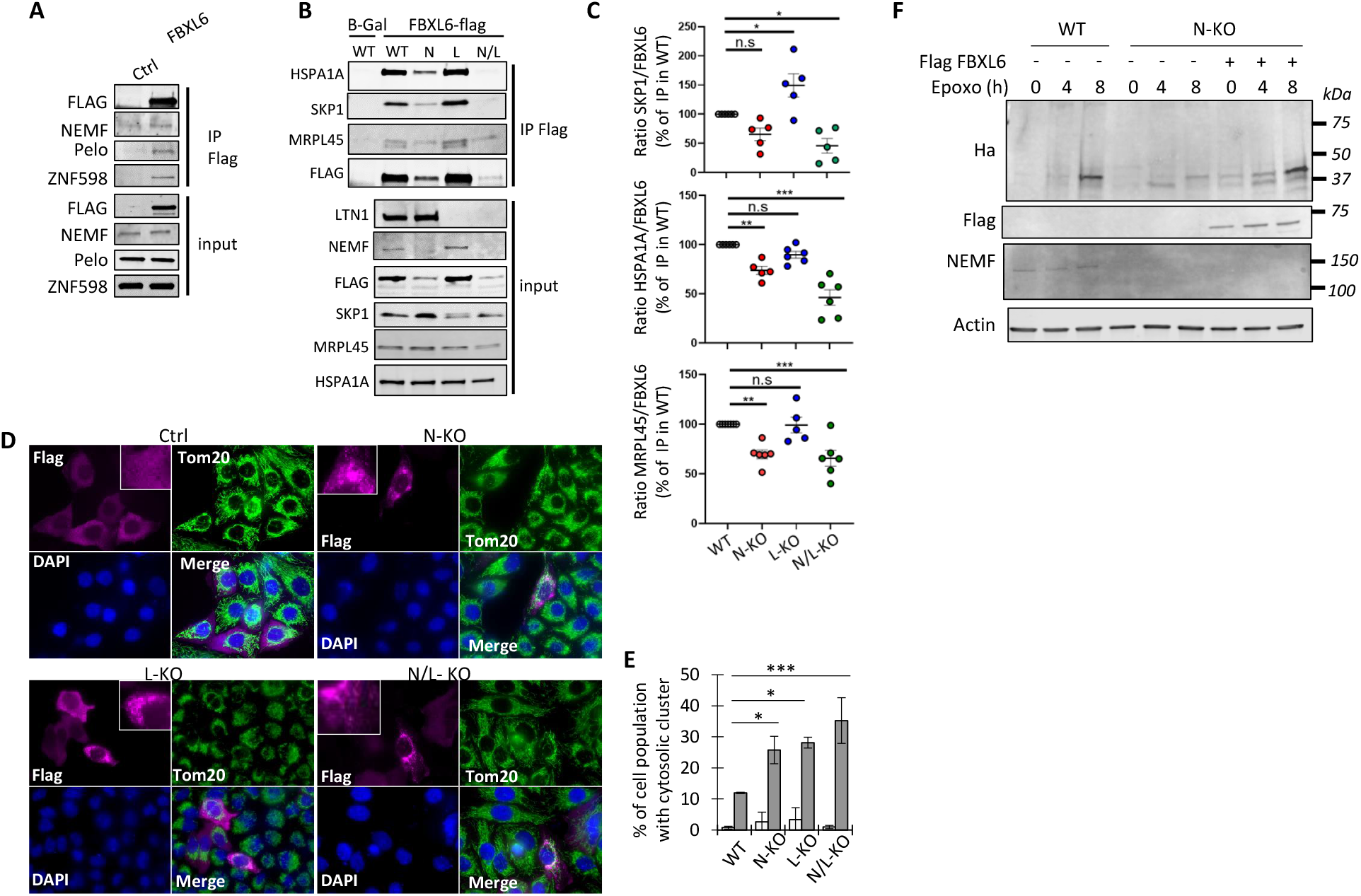
FBXL6 is involved in the RQC pathway. (A) Interaction of FBXL6 with RQC proteins were analyzed by Flag immunoprecipitation using cells transfected with FBXL6 flag or control (β-gal). Interaction with NEMF, PELO and ZNF598 were reveald by immunoblots using specific antibody. (B) Interaction of FBXL6-flag with partners and substrates were analyzed in LTN1-KO (L), NEMF-KO (N), double LTN1/NEMF KO (N/L) and control cells. FBXL6-Flag or GFP (β-gal) were expressed in these different cells and immunoprecipitations were performed using anti-FLAG and interactions with MRPL45, SKP1 and HSPA1A were analyzed by immunoblots. (C) Quantification of these interactions. Values represent protein levels normalized to Flag pulled down and expressed as percentage of immunoprecipation performed in control cells. (N =5-6, *** P < 0.001, ** P < 0.01, * P < 0.05, one way ANOVA, Holm-Sidak test) (D) Localization of FBXL6-flag was analyzed in control (ctrl), NEMF-KO (N-KO), LTN1-KO (L-KO) and NEMF/LTN1 double KO (N/L-KO) cells by immunofluorescence. Mitochondria and FBXL6 were labeled using anti-TOM20 (Green) and anti-flag (red) respectively. Nucleus was stained with Dapi (blue). (E) Quantification of cell number containing cytosolic FBXL6 (grey bars) or GFP (white bars) clusters. GFP distribution is represented in figure S6B. (n= 3, > 150 cells per n, * P < 0.05, *** P < 0.001, ordinary 1W ANOVA, Holm-Sidak test). (F) Control cells (WT) or NEMF-KO (N-KO) expressing MRPL45-HA-NS were transfected with FBXL6-Flag (or control) and treated epoxomicin for 0h, 4h or 8h. Relative accumulation of MRPL45 over the time was measured using anti-HA and normalized to actin (n=3).

As we reported above, FBXL6 interacts also with many chaperones including HSPA1A which played important role in quality control of nascent and newly synthesized proteins (Hartl and Hayer-Hartl 2002)(Tian et al. 2021) through physical interaction with the translating ribosome (Jaiswal et al. 2011). Thus, we analyzed the role of this chaperone in the FBXL6-dependent mechanism. Using blue native polyacrylamide gel electrophoresis (BN-PAGE) approach performed on whole cell extract, we observed build-up of MRPL45 in FBXL6-KO cells under the form of aggregates and these forms were absent in control cells (Fig. 6A-B). We found that these aggregates comigrated with HSPA1A. HSPA1A/MRPL45 aggregates were found at a lower molecular weight in FBXL4-KO cells confirming the involvement of FBXL4 in mitochondrial proteostasis whereas FBXL4 did not interact with MRPL45 (Fig. 6C). The different size of HSPA1A aggregates observed in FBXL4- and FBXL6-KO cells also pointed out discrepancies between related mechanisms. To test if these aggregates could be triggered by defective translation of mitochondrial proteins, we analyzed the subcellular localization of HSPA1A in presence of CAT-tailed mtGFP. Upon expression of control mtGFP, HSPA1A displayed diffused cellular localization (Fig. 6D) while expression of the CAT-tailed mtGFP induced the relocalization of HSPA1A to precise cytosolic foci (Fig. 6E). We found that expression of the CAT-tailed mtGFP led to the translocation of FBXL6 to these sites containing HSPA1 and the defective mitochondrial GFP (Fig. 6F-E, line scales). Increased colocalization between HSPA1A and FBXL6 in the presence of CAT-tailed was confirmed by high Pearson’s colocalization factor (Fig. 6F). Furthermore, immunoprecipitations of CAT-tailed versus WT MRPL45 revealed that HSPA1A had preferential interactions with the defective form of the ribosomal protein (Fig. 6G-H). However, HSPA1A/MRPL45-CAT interactions were not significantly modulated by the exogenous expression of either FBXL4 or FBXL6 (Fig. 6H). Finally, we found that silencing of HSPA1A dramatically reduced the ability of FBXL6 to bind MRPL45 (Fig.6 I-J).

**Figure 6.**
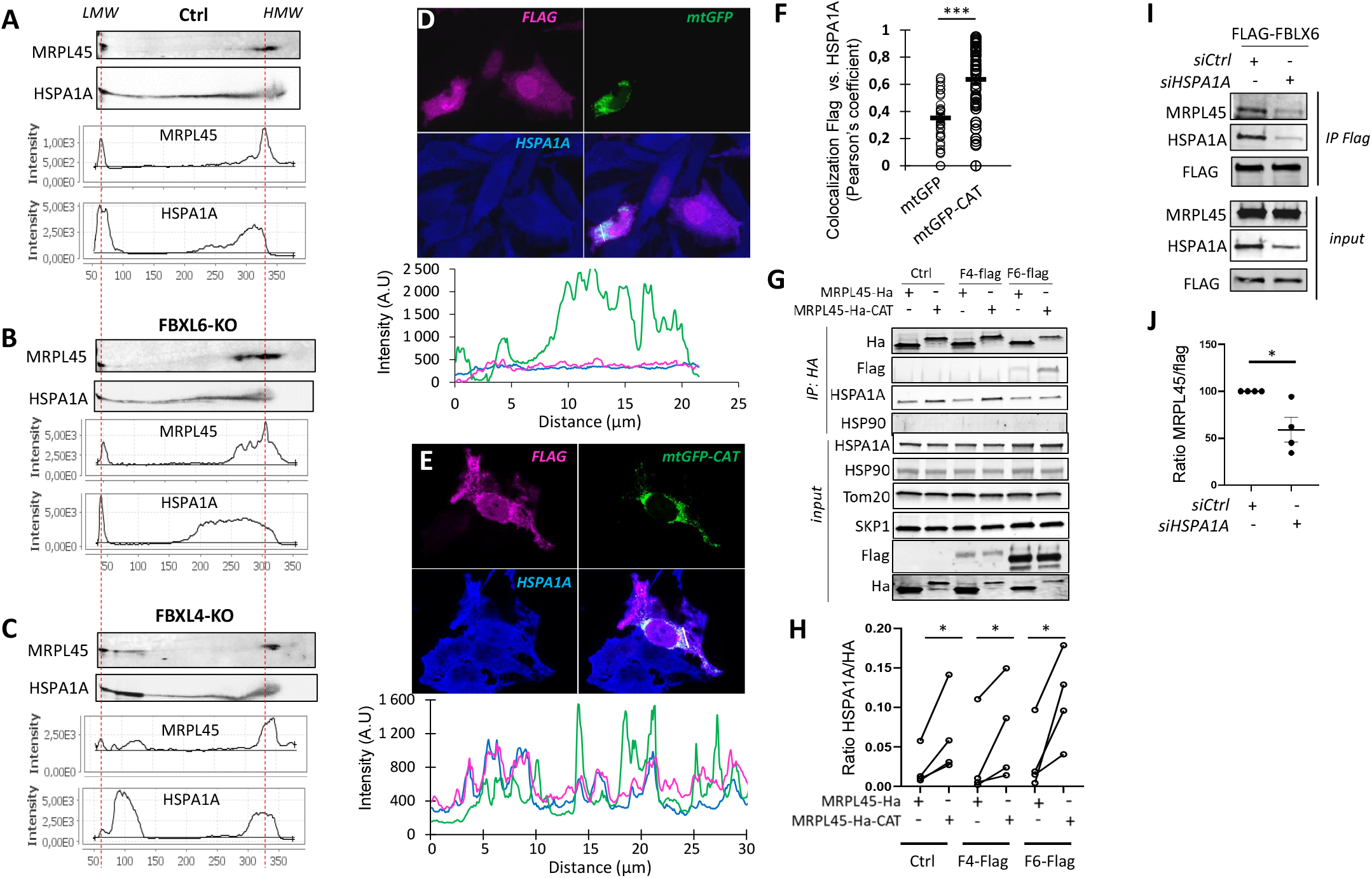
HSPA1A is involved in FBXL6-dependent mechanism. (A-C) Blue native polyacrylamide gel electrophoresis was performed on whole cell extracts from control cells (A), FBXL6-knockout (B) or FBXL4-KO (C) and control cells. Aggregates of mitochondrial ribosomal proteins and chaperones were studied using anti-mitochondrial ribosomal protein L45 (MRPL45) and anti-heat shock protein 1A1 (HSP1A1) antibodies. Plots displayed the quantification of the different profiles and red lines pointed out free MRPL45 at the low molecular weight side (LMW) and the native mitochondrial ribosome at the high molecular weight side (HMW). (D-E) Colocalization of FBXL6-Flag and HSPA1A was assayed by immunofluoresence in presence of WT mitochondrial GFP (mitoGFP; D) or C-terminal alanine and threonine (CAT)-tailed mitochondrial GFP (mtGFP CAT; B). FBXL6 and heat shock protein A1A (HSPA1A) were labeled with anti-Flag (red) and anti-HSPA1A (blue). Linescales show the recruitment of FBXL6 and HSPA1A to mtGFP-CAT aggregates. (F) HSPA1A and FBXL6 colocalization was quantified by measuring the Pearson’s correlation coefficient in presence of mtGFP or mtGFP CAT. Each dot is the value for one cell (N = 3, *** P < 0.001, unpaired T-test). (G-H) Cells expressing FBXL4-Flag or FBXL6, or control cells were transfected with WT MRPL45- or MRPL45-HA-CAT. HA or Flag-immunoprecipitation was performed and the immunoprecipitated proteins were analyzed using immunoblots (I). Quantification of HSPA1A pulled down with HA expressed as HSPA1A/HA band ratio (J; N = 4, * P < 0.05, paired t-test). (I-J) HSPA1A was downregulation by using siRNA in cells expressing FBXL6-Flag and Flag immunoprecipiation was performed (G). Binding of MRPL45 was analyzed by immunoblots and relative level was normalized to flag (H). (N = 4, * P < 0.05, unpaired T-test).

Taken together, these results demonstrate that HSPA1A carries defective ribosomal proteins to FBXL6.

## Discussion

In the current study, we show that FBXL6 is a cytosolic E3 ubiquitin ligase that contributes to the mitochondrial proteostasis. FBXL6 acts at the interface between the cytosolic translation and chaperones to ensure the quality control of newly synthesized mitochondrial ribosomal proteins. Cells carrying deletion of FBXL6 reduced degradation rates of defective mitochondrial ribosomal protein and altered mitochondrial metabolic activities.

### FBXL6, an element of the mitoRCQ in mammalian cells

Most mitochondrial proteins are translated in the cytosol under the form of precursors which are then transported to the different mitochondrial compartments. This synthesis represents also two successive sites implicated in the quality control of these proteins. At the level of the organelle, the translocase of the outer membrane (TOM) is the receptor of key proteins involved the quality control such as PINK1 which triggers a global quality control by mitophagy (Lazarou et al. 2012) or molecular Chaperones Hsp90 and Hsp70 which are involved in the quality control of newly synthesized proteins (Young, Hoogenraad, and Hartl 2005). Accordingly, blockade of mitochondrial protein import triggers cellular stress responses including the upregulation of the unfolded protein response and heat shock proteins (HSP) (Boos et al. 2019)(Wrobel et al. 2015). Notably, this process prevents incorporation of defective proteins as revealed by Mohanraj et al. who showed that the mutated mitochondrial protein COA7 has a slower mitochondrial import rate and that mislocalized proteins are degraded in the cytosol by the proteasome (Mohanraj et al. 2019). In the cytosolic side, ribosomes represent a major platform that orchestrates mRNA and nascent chain quality control by sensing the mRNA integrity, by recruiting folding proteins, and by supporting the ubiquitination of defective proteins, (Pechmann, Willmund, and Frydman 2013). By demonstrating the role played by Vms1 at the interface between the RQC and the mitochondrial import, Izawa and colleagues revealed a straddling mechanism, the mitoRQC that unified the cytosolic translation to mitochondrial protein quality control in yeast (Izawa et al. 2017). Complementing these findings it was shown that Ubx2, which also participates in the ERAD process, directly interacts with the TOM complex and recruits Cdc48 to tag defective newly synthesized proteins for proteasomal degradation (Mårtensson et al. 2019). Such a mechanism has not yet been described in higher eukaryotes but this existence is highly probable. In this line, Wu et al. have demonstrated in *Drosophila* that mitochondrial damages induced ribosome stalling at the OMM which promote the recruitment of PELO and ABCE1 at these sites (Wu et al. 2018). In our current study, we detailed the role of the E3 ubiquitin ligase FBXL6 in the quality control of neosynthesized mitochondrial proteins in human cells. We found that FBXL6 binds to ZNF598, PELO and NEMF, keys actors of the RQC. Furthermore, FBXL6 interacts with several chaperones and co-chaperones involved in the folding and transport of newly synthesized polypeptides, such as HSPA1A, HSPA4, and HSP90AB1, and proteins involved in the sorting and delivering of nascent/neosynthesized proteins at the OMM (HSPA8, HSP90AA1,) (Young, Hoogenraad, and Hartl 2005). We identified that newly synthesized mitochondrial ribosomal proteins are substrates of this enzyme. Thus, since FBXL6 is not located to the mitochondria, interactions with mitochondrial ribosomal proteins occur with the cytosolic precursors. Cells lacking FBXL6 failed to eliminate defective mitochondrial proteins resulting from altered translation while the expression of CAT-tailed and non-stop mitochondrial ribosomal proteins was sufficient to concentrate this E3 ubiquitin ligase to precise cytosolic foci that contained defective proteins and chaperones. According to these data, the FBXL6-dependent mechanism takes place at the interface between the OMM and the ribosome quality control hub comprising RQC proteins and chaperones (Fig. 7). Thus, this mechanism confirms the existence of a mitoRQC-like process in mammalian cells

**Figure 7.**
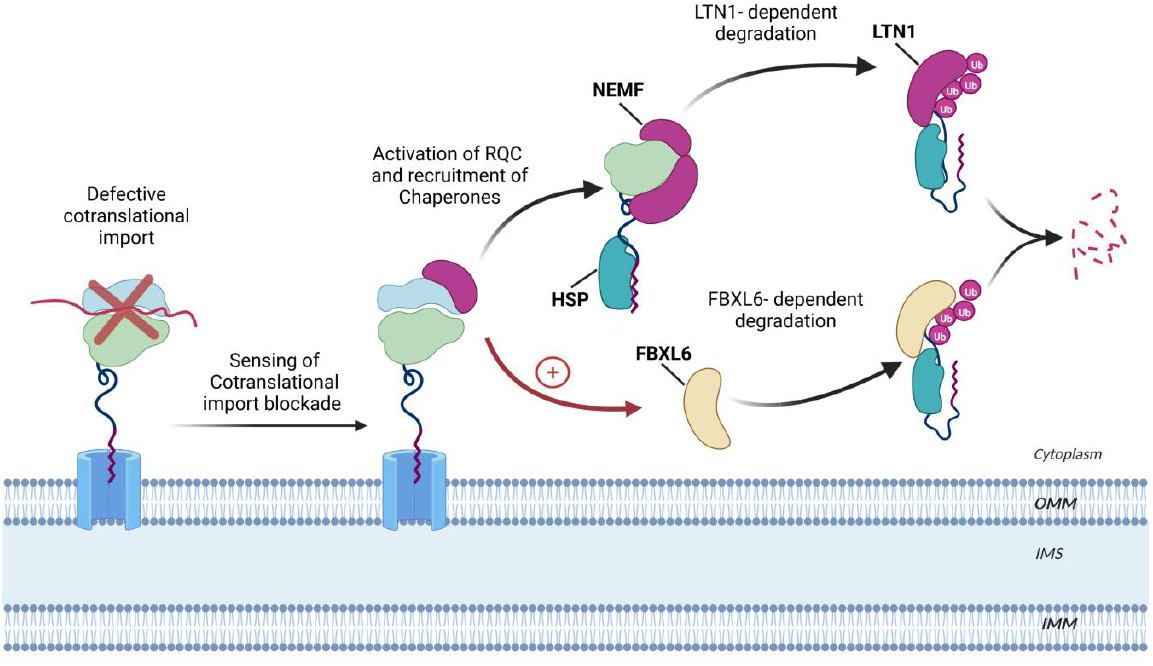
Model of FBXL6-dependent mechanism. Activation of the RQC due to defective translation of mitochondrial proteins triggers FBXL6-dependent mechanism. In this mechanism, HSP are required to carry defective ribosomal proteins for ubiquitination and this FBXL6-dependent mechanism works in parallel of LTN1-dependent degradation.

### Implication of E3 ubiquitin ligases in mitoRCQ

Notably, our findings rise also questions regarding the substrate specificity during the RQC process. Uses of GFP-based reporters have allowed to identify and to understand the RQC molecular mechanism but these approaches provided a limited comprehension about substrate specificity. Accordingly, it is unknown if proteins with different physical properties (e.g. folding stability, hybrophobicity, leading sequence etc…) are similarly recognized and managed by the RQC machinery or if some group of proteins (e.g ribosomal proteins) require a specific response/machinery. E3s ligases are enzymes that can provide a certain elasticity to the RQC response. In our study, we used modified endogenous proteins and we showed that FBXL6 targets specifically defective mitochondrial ribosomal proteins while other defective mitochondrial proteins (i.e. SDHA nor UQCRC2) are not impacted. Therefore, these results support the existence of certain specificity between defective substrate and E3 at least in mammalian cells. In yeast, Duttler et al. showed that ubiquitination during translation affects up to 5% of nascent proteins and they demonstrated that these modifications are carried out by several E3 ubiquitin ligases (Duttler, Pechmann, and Frydman 2013). Some of them promote the degradation of non-stop protein (Ltn1) or nascent unfolded proteins (Hul5, Hrd1). Recently, the group of Joazeiro revealed that CRL2^KLHDC10^ and Pirh2 participate to the RQC in mammalian cells and can eliminate defective proteins in absence of LTN1 (Thrun et al. 2021) (Bengtson and Joazeiro 2010). They proposed that the CAT tail serves a degron signal.

Beside FBXL6, it is likely that other E3 ubiquitin ligases participate to the mitoRCQ and this participation might depend on the physiological context like the preferential recognition of substrates, the responses to physiological stimuli or tissue specificity. Notably, MARCH5, an OMM located E3 ubiquitin ligase previously reported to regulate mitochondrial morphology, orchestrates the cytosolic retention of nascent/neosynthesized proteins by promoting their ubiquitination (Karbowski, Neutzner, and Youle 2007)(Phu et al. 2020). Another E3 ubiquitin ligase, NOT4, seems to have an important role in the mitoRQC process because Wu et al. have revealed that NOT4 ubiquitinates ABCE1, a protein involved in the splitting of the ribosomal subunits, upon aberrant translation (Wu et al. 2018). These authors showed that the collapse of the mitochondrial membrane potential induces cytosolic ribosome stalling and that the accumulation of ubiquitinated ABCE1 triggers mitophagy mediated by the PINK1 pathway.

FBXL4, another member of the FBXL family, might be also involved in the mitoRQC by targeting exclusively mitochondrial proteins. Mutations of FBXL4 is responsible for a severe mitochondrial disease characterized by early-onset lactic acidemia, hypotonia, and developmental delay caused by severe encephalomyopathy (Gai et al. 2013)(Bonnen et al. 2013)(Huemer et al. 2015). The molecular mechanism responsible for this disease remains uncharted. By summarizing clinical data collected from 87 patients with FBXL4 mutations, El-Hattab and colleagues showed that 92% of these patients displayed a depletion of mtDNA in muscle ranging between 10 and 60%. Notably, 30% of these patients did not feature a decrease in OXPHOS enzymatic activities in muscle tissue (El-Hattab et al. 2017). Other studies have also pointed out that the mitochondrial morphology is altered in patient-derived fibroblasts exhibiting shortened and highly fragmented mitochondria (Sabouny et al. 2019)(Gai et al. 2013). In addition, Larsson’s group recently showed that FBXL4 loss of function induces activation of mitophagy and this activation can be responsible for mtDNA and mitochondrial protein depletion in an FBXL4 KO mouse model (Alsina et al. 2020). Overall, FBXL4 loss of function leads to disparate alterations of the organelle including impaired metabolism, altered morphology, mitophagy, defective mtDNA maintenance. To date, it remains unknown how the loss of the FBXL4 E3 ubiquitin ligase activity triggers these mitochondrial alterations. Implications of FBXL4 in the mitoRQC can explain most of disparate features observed in patients’ cells. Indeed, we observed strong similarities between the FBXL4- and FBXL6-dependent mechanisms. First, deletion of both FBXL4 and FBXL6 inhibits the G2/M transition, the protein synthesis phase and this inhibition occurs only when the mitochondrial metabolism is required to produce energy. Like FBXL6, FBXL4 physically interacts with chaperones such as HSPA1A and deletion of the E3 induces aggregation of mitochondrial proteins in complexed with HSPA1A. Moreover, lack of FBXL4 inhibits the degradation of CAT-tailed mitochondrial proteins showing that the FBXL4-dependent mechanism is also linked to the quality control of defective proteins. The strict mitochondrial localization of FBXL4 suggests that this protein ensures the exclusively quality control of mitochondrial proteins. Thus, it could be expected that the metabolic dysfunctions, increased mitochondrial damages, mitochondrial fragmentation and mitophagic degradation reported in FBXL4-KO and FBXL4 patient’s’ cells result from incremental aggregation of the proteins in the mitochondrial membranes. Notably, it was recently proposed that loss-of-function mutations in *ANKZF1* are responsible infantile-onset inflammatory bowel disease and patients’ cells carrying these mutations exhibit deregulation of mitochondrial integrity (Van Haaften-Visser et al. 2017).

## Author contributions

J.L. and G.B. conceived the study. J.L., W.M., H. R., and G.B. performed the biochemical experiments. J.L. designed and conceived genetic constructs. J-W. D. performed the mass spectrometry analyses. C.L. performed the activities and oxygraphy assays. W.M. and H. R., performed the cell cycle assays. P. N. and AM Duchene have generated HCT116 TurboID cells and performed proximity labelling assay. G.B. wrote the manuscript with contributions of all authors.

## Acknowledgments.

We thank Dr Claudio Joazeiro for LTN1, NEMF and LTN1/NEMF KO cells. Giovanni Bénard was supported by “agence national de la recherche”, ANR, (n°ANR-17-CE14-0039-01), by Region Nouvelle-Aquitaine (2019-1R30124) and by AFM. RP Ngondo were supported by an ANR grant (ANR-18-CE12-0021-01 “Polyglot”) and by the French National Program “Investissement d’Avenir” (Labex MitoCross). We thank A. Sequiera for technical supports. We acknowledge, Johana Chicher from the Plateforme Protéomique Strasbourg Esplanade (CNRS) for performing the TurboID mass-spectrometry analysis.

---

Figure S1.

(A-B) FBXL4 relative mRNA (A) or protein (B) expression was measured in 3 different FBLX4 KO clonal cell populations. Protein and mRNA relative expression was measured by western blot and qPCR, respectively. F4-1 clone was deleted of one allele and F4-2 and F4-3 are full KO clones. For Immunoblot, we used our home made anti-FBXL4 (thermoScientific customed antibody) confirmed through FBLX4-Flag overexpression in HEK.

(C-D) FBXL6 relative mRNA (A) or protein (D) expression was measured in 3 different FBLX4 KO clonal cell populations. Protein and mRNA relative expression was measured by western blot and qPCR, respectively. All FBXL6 clones are full KO.

(E) FBXL6-KO cells were transfected with FBXL6 or control (GFP). After 24h transfection, cells were switched to oxidative media for 48h of incubation. Each dot represents number of cell after treatment and blacked dashed line is the number of seeded cells. (N= 4, *p<0.05, unpaired T-test)

(F) Inhibition of oxygen consumption rate in the different FBXL4-KO and FBXL6-KO cells as compared to control. Values are expressed as percentage of inhibition of OCR measured in control cells (N= 8) (G-H) Relative expression of mitochondrial proteins was measured by immunoblots performed on total cell extract of three different FBXL6-KO hela clones (G). Expressions were normalized to actin and expressed as % of control cell (H). (n= 3-9, ANOVA Kruskal-Wallis test, * p<0.05, **p<0.01, *** p<0.001).

Figure S2. Analyses of mitochondrial localization of FBXL4 and FBXL6 using mitominer (Smith and Robinson 2016).

Figure S3.

(A) Immunoprecipitation assays performed on HEK cell expressing β-gal (ctrl), FBXL3 or FBXL6. Interaction with MRPL45 and SKP1 was analyzed by immunoblots.

(B-C) Localization and expression of V5 tagged FBXL6-TurboID stably expressed in HCT116 cells were analyzed by immunofluorescence (B) and immunoblot (C). V5 tagged FBXL6-TurboID was detected using using V5 immunostaining. HCT116 cells stably expresing V5 tagged GFP-TurboID were used as control.

(D) Immunoblot showing levels of protein biotinylation in HCT116 cells expressing GFP-TurboID or FBXL6-TurboID after 0h, 0.5h or 16h incubation with biotin. Biotinylated proteins and TurboID chimeric proteins were revealed using streptavidin-coupled to HRP and anti-V5, respectively.

(E-F) Biotinylated proteins were identified using mass spectrometry after streptavidin pull down. Plots showing the fold changes of biotinylated protein levels and p-values in FBXL6-TurboID- and GFP-TurboID-expressing cells. N = 3.

(G) Gene ontology analyses of proteins specifically interacting with FBXL6 at 30min (G). Mitochondrial pathways are underlined.

Figure S4.

(A) Immunoblots showing migration profile of WT and CAT-tailed cytosolic green fluorescent protein (cGFP and cGFP-CAT) and WT and CAT-tailed mitochondrial green fluorescent protein (mtGFP and mtGFP-CAT). GFP expression were measured using anti GFP. Cells were treated with epoxomicin or MG132 for 8h.

(B-C) Immunofluorescence performed on Hela cells expressing cGFP and cGFP-CAT (B) or mtGFP and mtGFP-CAT (C). Immunofluorescence were performed using anti-TOM20 (red) and GFP staining on cells treated with epoxomicin and vehicle for 8h. As compared to cells expressing GFP, aggregates were observed in cGFP-CAT transfected cells, which were enhanced in the presence of epoxomicin.

Figure S5.

(A-B) FBXL6-KO and control cells were transfected with MRPL42-HA (A) or MRPL42-HA-CAT (B) and treated with cycloheximide for 0h, 2h 4h and 8h. The protein levels of MRPL42-HA and MRPL42-HA-CAT were evaluated using anti-Ha and immunoblots. Curves correspond to the protein quantification normalized to protein level at t = 0 (N = 3).

(C) FBXL6-KO, FBXL4-KO and control cells were transfected with SDHA-HA-CAT and treated with cycloheximide for 0h, 2h 4h and 8h. The protein levels were evaluated using immunoblots and anti HA. Curves correspond to the protein quantification normalized to protein level at t = 0 (N = 3).

(D) Degradation rate of MRPL45-HA-NS in WT or FBXL-KO cells. Cells were treated with cycloheximide and relative expression of MRPL45-HA-NS was measured by immunoblot using anti-Ha. Asterisk is unspecific band. Curves correspond to the protein quantification normalized to protein level at t = 0 (N = 4).

Figure S6.

(A) Expression of MRPL45 was analyzed in FBXL6-KO (red) and control cells (blue) transfected with siRNA against NEMF, LTN1 ou Luciferase (ctrl). (N = 3, * p<0.05, ns; non significant, paired t-test)

(B) HEK cells expressing flag-FBXL6 were transfected PELO, NEMF ZFN598 and LTN1 siRNA. Flag immunoprecipitation was performed and binding of SKP1, HSPA1A and MRPL45 was analyzed by immunoblots

(B) Distribution of GFP (green) in control (ctrl), NEMF-KO (N-KO), LTN1-KO (L-KO) and NEMF/LTN1 double KO (N/L-KO) cells was evaluated by immunofluorescence. Mitochondria and nucleus were labeled using anti-TOM20 (red) and DAPI, respectively.

**Table S1:** excel proteomic KO vs ctrl

**Table S2:** excel proteomic IP ctr vs FBXL

**Table S3:** excel proteomic IP ctr vs FBXL GO

**Table S4:** Excel proteomic turboID ctrl vs FBXL

## STAR Methods

### Resource table

| REAGENT or RESOURCE | SOURCE | IDENTIFIER |
| --- | --- | --- |
| **Antibodies** |  |  |
| Mouse Anti-ATP5A mAb | Santa Cruz Biotech | Cat# sc-136178, RRID:AB_2061764 |
| Mouse anti-beta-Actin mAb | Sigma Aldrich | Cat# A1978, RRID:AB_476692 |
| Mouse anti-COX2 mAb | Santa Cruz Biotech | Cat# sc-514489 |
| Mouse anti-COX4 mAb | Santa Cruz Biotech | Cat# sc-376731 |
| Mouse anti-FLAG M2 mAb | Sigma-Aldrich | Cat# F3165, RRID:AB_259529 |
| Rabbit anti-FBXL4 pAb | Invitrogen | Custom antibody AB2862 |
| Rabbit anti-FBXL6 pAb | Invitrogen | Custom antibody AB2863/AB2864 |
| Mouse anti-GFP mAb | Roche | Cat# 11814460001, RRID:AB_390913 |
| Rabbit anti-GAPDH pAb | Santa Cruz Biotech | Cat# sc-25778, RRID:AB_10167668 |
| Rabbit anti-HA mAb | Cell Signalling | Cat# 3724, RRID:AB_1549585 |
| Mouse anti-HSP60 | Novus | Cat# NBP2-22440 |
| Rabbit anti-HSP90 mAb | Cell Signalling | Cat# 4877, RRID:AB_2233307 |
| Rabbit anti-HSPA1A pAb | ThermoFisher | Cat# PA5-28003, RRID:AB_2545479 |
| Rabbit anti-RNF160 pAb | Proteintech | Cat# PR-28452-1-AP |
| Rabbit anti-MRPL42 | Novus | Cat# NBP1-83171, RRID:AB_11006589 |
| Rabbit anti-MRPL45 | Novus | Cat# NBP1-82764, RRID:AB_11019553 |
| Mouse anti-MRPS34 | Novus | Cat# NBP2-45432 |
| Mouse anti-Myc tag mAb | Merck Millipore | Cat# 05-419, RRID:AB_309725 |
| Rabbit Anti-Nemf | Bethyl Laboratories | Cat# A305-758A-M |
| Mouse anti-NDUFS3 mAb | Santa Cruz Biotech | Cat# sc-374282, RRID:AB_10991315 |
| Mouse anti-OPA-1 mAb | BD Biosciences | Cat# 612607, RRID:AB_399889 |
| Rabbit anti PELO | GeneTex | Cat# GTX120446 |
| Rabbit anti-RPL9 | Novus | Cat# NBP1-82853, RRID:AB_11027189 |
| Mouse anti-SDHA mAb | Santa Cruz Biotech | Cat# sc-166909, RRID:AB_10611174 |
| Mouse anti-SKP1 | Novus | Cat# NBP2-37539 |
| Rabbit anti-Tom20 (FL-145) pAb | Santa Cruz Biotech | Cat# sc-11415, RRID:AB_2207533 |
| Rabbit anti-UQCRC2 pAb | Proteintech | Cat# 14742-1-AP, RRID:AB_2241442 |
| Mouse anti-V5-Tag mAb | Biorad | Cat# MCA1360, RRID:AB_322378 |
| Rabbit anti-ZNF598 | GeneTex | Cat# GTX119245 |
| IRDye 800CW Donkey anti-Mouse IgG antibody | LI-COR Biosciences | Cat# 926-32212, RRID:AB_621847 |
| IRDye 800CW Goat anti-Rabbit IgG antibody | LI-COR Biosciences | Cat# 926-32211, RRID:AB_621843 |
| IRDye 680RD Donkey anti-Mouse IgG antibody | LI-COR Biosciences | Cat# 926-68072, RRID:AB_10953628 |
| IRDye 680RD Donkey anti-Rabbit IgG antibody | LI-COR Biosciences | Cat# 926-68073, RRID:AB_10954442 |
| Goat Anti-Mouse IgG (H L)-HRP Conjugate antibody | Bio-Rad | Cat# 170-6516, RRID:AB_11125547 |
| Goat Anti-Rabbit IgG (H L)-HRP Conjugate antibody | Bio-Rad | Cat# 170-6515, RRID:AB_11125142 |
| Goat anti-Mouse IgG (H+L) Cross-Adsorbed Secondary Antibody, Alexa Fluor 350 | Thermo Fisher Scientific | Cat# A-11045, RRID:AB_2534100 |
| Goat anti-Rabbit IgG (H+L) Cross-Adsorbed Secondary Antibody, Alexa Fluor 350 | Thermo Fisher Scientific | Cat# A-11046, RRID:AB_2534101 |
| Goat anti-Mouse IgG (H+L) Cross-Adsorbed Secondary Antibody, Alexa Fluor 488 | Thermo Fisher Scientific | Cat# A-11001, RRID:AB_2534069 |
| Goat Anti-Rabbit IgG (H+L) Antibody, Alexa Fluor 488 Conjugated | Thermo Fisher Scientific | Cat# A-11008, RRID:AB_143165 |
| Goat anti-Mouse IgG (H+L) Cross-Adsorbed Secondary Antibody, Alexa Fluor 568 | Thermo Fisher Scientific | Cat# A-11004, RRID:AB_2534072 |
| Goat anti-Rabbit IgG (H+L) Cross-Adsorbed Secondary Antibody, Alexa Fluor 568 | Thermo Fisher Scientific | Cat# A-11011, RRID:AB_143157 |
| Goat anti-Mouse IgG (H+L) Cross-Adsorbed Secondary Antibody, Alexa Fluor 647 | Thermo Fisher Scientific | Cat# A-21235, RRID:AB_2535804 |
| Goat anti-Rabbit IgG (H+L) Cross-Adsorbed Secondary Antibody, Alexa Fluor 647 | Thermo Fisher Scientific | Cat# A-21244, RRID:AB_2535812 |
| Streptavidin Mag Sepharose® | Cytiva | Cat# 28-9857-38 |
| Pierce™ Anti-HA Magnetic Beads | Thermo Fisher | Cat# 88836 |
| Anti-FLAG® M2 Magnetic Beads | Sigma Aldrich | Cat# M8823 |
| **Chemicals and kits** |  |  |
| Epoxomycin | UBPBIO | Cat# F1400-5 |
| Cycloheximide | Sigma-Aldrich | Cat# 01810 |
| MG132 | UBPBIO | Cat# F1101 |
| Dithiothreitol | Sigma Aldrich | Cat# D5545 |
| Protease Inhibitor Cocktail | Sigma-Aldrich | Cat# P8340 |
| Bovine Serum Albumin | Sigma-Aldrich | Cat# A7906-500G |
| 2-Mercaptoethanol | Sigma Aldrich | Cat# M3148 |
| Digitonin | Sigma Aldrich | Cat# D141-100MG |
| Carfilzomib (PR-171) | UBPBio | Cat# F1300 |
| Pierce™ BCA Protein Assay Kit | Thermo Scientific | Cat# 23225 |
| Adenosine 5'-triphosphate disodium salt hydrate | Sigma Aldrich | Cat# A7699-5G |
| CellTiter-Glo® 2.0 Cell Viability Assay | Promega | Cat# G9242 |
| **Experimental Models: Cell Lines** |  |  |
| Wild-type HEK293 | ATCC | Cat# CRL-1573;<br>RRID: CVCL_0045 |
| Wild-type HeLa cells | Sigma Aldrich | ECACC Cat# 93021013,<br>RRID:CVCL_0030 |
| FBXL4 KO HeLa stable | This manuscript | N/A |
| FBXL6 KO HeLa stable | This manuscript | N/A |
| HCT 116 cell line | ATCC | ATCC, CCL-247 |
| TurboID FBXL4 HCT116 stable | This manuscript | N/A |
| TurboID FBXL6 HCT116 stable | This manuscript | N/A |
| **Recombinant DNA** |  |  |
| pCMV-beta-galactosidase | Gift | N/A |
| pCMV6-AC-GFP | Origene | Cat#: PS100010 |
| FBXL3-Myc | Genscript | Clone ID: OHu06084C |
| FBXL4-Myc-DDK | Origene | Cat# RC208817 |
| FBXL6-Myc-DDK | This paper | N/A |
| mlsFBXL4-GFP | This paper | N/A |
| mlsFBXL6-GFP | This paper | N/A |
| mls-HA-ubi | (Lavie et al. 2018) | N/A |
| Ha-ubi | (Lavie et al. 2018) | N/A |
| Cyto-GFP | This paper | N/A |
| Cyto-GFP-CAT | This paper | N/A |
| Mito-GFP | This paper | N/A |
| Mito-GFP-CAT | This paper | N/A |
| MRPL45-HA-CAT | This paper | N/A |
| MRPL45-HA-NS | This paper | N/A |
| **Other** |  |  |
| Fugene HD Transfection Reagent | Promega | Cat# E2311 |
| RIPA buffer | Sigma-Aldrich | Cat# R0278 |
| Laemmli Sample Buffer 4x | Bio-Rad | Cat# 1610747 |
| Laemmli Sample Buffer 2x | Sigma Aldrich | Cat# S3401 |
| Precision Plus Protein Dual Color Standards | Bio-Rad | Cat# 1610374 |
| 10x Tris/Glycine/SDS | Bio-Rad | Cat# 1610772 |
| Mini-PROTEAN TGX Gels | Bio-Rad | Cat# 4561093 |
| Mini-PROTEAN TGX gels IPG | Bio-Rad | Cat# 4561091 |
| Trans-Blot Turbo RTA Midi 0.2 µm PVDF Transfer Kit | Bio-Rad | Cat# 1704273 |
| Penicillin-Streptomycin | ThermoFisher Scientific | Cat# 15140122 |
| DMEM, high glucose, GlutaMAX™ Supplement, pyruvate | ThermoFisher Scientific | Cat# 31966021 |
| DMEM, no glucose | ThermoFisher Scientific | Cat# 11966025 |
| HEPES (1M) | ThermoFisher Scientific | Cat# 15630056 |
| L-Glutamine (200 mM) | ThermoFisher Scientific | Cat# 25030024 |
| Sodium Pyruvate (100 mM) | ThermoFisher Scientific | Cat# 11360070 |
| D-(+)-Galactose | Sigma Aldrich | Cat# G0625 |
| Trypsin-EDTA (0.25%), phenol red | ThermoFisher Scientific | Cat# 25200072 |
| SuperSignal™ West Pico PLUS | Thermo Scientific | Cat# 34577 |
| SuperSignal™ West Femto | Thermo Scientific | Cat# 34095 |
| Trypsin-EDTA (0.05%) | Gibco | Cat# 25300062 |
| 2-Mercaptoethanol | Sigma Aldrich | Cat# M3148 |
| NativePAGE 20x running buffer | Invitrogen | Cat# BN2001 |

### CONTACT FOR REAGENT AND RESOURCE SHARING

Further information and requests for resources and reagents should be directed to and will be fulfilled by the Lead Contact, Giovanni Bénard.

### EXPERIMENTAL MODEL AND METHOD DETAILS

#### Cell lines, culture conditions, and transfections

Hela were obtained from Merck and certified by the European Collection of Authenticated Cell Cultures (ECACC) and HEK cells from ATCC(CRL-1573). HCT116 cells were purchased from ATCC (CCL-247, ATCC). Cells were cultured in glucose media consisting of Dulbecco’s modified Eagle medium (DMEM) medium containing 25 mM glucose. All media were supplemented with 10% heat-inactivated fetal bovine serum, 1 mM sodium pyruvate, MEM non-essential amino acids, 100 U/mL penicillin, and 100 μg/mL streptomycin. Cells were cultured in a 5% CO_2_ atmosphere at 37 °C and transfected using FugeneHD (Roche, Basel Switzerland) according to the manufacturer’s protocol.

For experiments in the presence of different energy-related substrates, high-glucose medium was removed and replaced with DMEM glucose-free medium containing 10 mM galactose and supplemented with either 4 mM glutamine (glutamine media) or 5 mM glucose (glucose media). Cells were cultured for 48 h prior to the experiments.

For protein turnover assays, cells were treated with 10 μM epoxomicin and 50 μg/mL cycloheximide (final concentration).

#### In vitro cell growth assay

Cell growth studies were carried out by plating 1.7 × 10^4^ cells in a 24-well plate (Corning) containing 1 ml of glucose medium in triplicates. After 24hrs (Day 0), the medium was removed and replaced with fresh glucose medium or galactose medium. The galactose medium consisted of DMEM deprived of glucose (no. 11966-025; Life Technologies, Inc.), supplemented with 10 mm galactose, 2 mm glutamine (4 mm final), 10 mm HEPES, 1 mm sodium pyruvate, and 10% FCS. HeLa cells were kept in CO_2_ 10% at 37°C or galactose medium. At daily intervals, cells were harvested by trypsinization and counted using a hemocytometer. For rescue experiments, FBXL6 KO cells (clone) were seeded at 4 × 10^4^. Cells were transfected with FBXL6 or GFP and counted after 48h incubation with galactose media.

#### Generation of control, FBXL4-, and FBXL6-KO cells

FBLX4 and FBXL6 KO Hela cells were generated in our CRISPR facility (CRISP’EDIT University of Bordeaux, France). FBXL4 KO and FBXL6 KO HeLa cells were generated by CRISPR in wild-type HeLa cells (ATCC, ECACC 93021013). SpCas9 target sequence crRNA1-FBXL4 = Hs.Cas9.FBXL4.1.AA: CCCCACAAATCTTATACGAC and crRNA2-FBXL4 = Hs.Cas9.FBXL4.1.AB: TGGGAAGCATTCACCTTCGT located in the exon 5 for FBXL4 and crRNA1-FBXL6 = Hs.Cas9.FBXL6.1.AA TCAGCCACAACTTGCGCATT and crRNA2-FBXL6 = Hs.Cas9.FBXL6.1.AB located in the exon 5 were designed using a CRISPOR algorithm. Single clones were then selected and PCR amplification with subsequent Sanger sequencing of the targeted FBXL4 or FBXL6 sequence were performed according the supplier’s recommendations with external specific primers for FBXL4 (5’-GTGACATATTGGGCATGGGTCTAT-3’ and 5’-CCGAGGACAGCACTGCTAAAC-3’) and for FBXL6 (5’-TAGCCTGGACCTACAGCACT-3’ and 5’-CTGAAGGGGAATGCTA-3’).

#### Generation of GFP, FBXL4, and FBXL6 turboID cells

Stable cell lines expressing TurboID-fusion proteins were generated by transducing HCT116 (CCL-247, ATCC) cells with lentiviral supernatant produced following the transfection (Lipofectamine 2000, Invitrogen) of 293T cells with packaging plasmids (pVSV and psPAX5) together with the pNG114 (GFP-TurboID), pNG115 (TurboID-FBXL4) or pNG116 (TurboID-FBXL6) vectors. The cells are grown at 37°C, 5% CO_2_ and maintained in DMEM (Sigma) with 10% Fetal bovine serum (Gibco) and 1% Pen/strep (Gibco). Transduced cells were selected with 2μg/mL of puromycin (Invitrogene) for at least one week.

#### Analysis of gene expression by quantitative PCR

Total RNAs were extracted from cells using the RNeasy Plus Mini kit (Qiagen) following the manufacturer’s recommendations. First strand cDNA was synthesized in a 20 μl volume using 1 μg of total RNA and iScript™ Reverse Transcription Supermix (Bio-Rad). PCR reactions were carried out in 25 μl volumes using SYBR Green PCR master mix (Bio-Rad) and 0.2 μM specific primers The following primer sets were deployed for real-time PCR analysis: h-FBXL4-F (TCGGGGAAGGGCCAAATAAT) and h-FBXL4-R (TTGCCCAGTATGGTTGCAGA), h-FBXL6-F (GCCAGGGGTTCAGTGAGAAG) and h-FBXL6-R (CAGGTTGAGGTAGAGCAGGC), h-RPLP0-F (GGCGACCTGGAAGTCCAACT) and h-RPLP0-R (CCATCAGCACCACAGCCTTC), h-GUSB-F (GGAGAGCTCATTTGGAATTTTGCCG) and h-GUSB-R (TGGCTACTGAGTGGGGATACCTGG), m-FBXL4-F (CCAGCTCCCAAACTTGCAGA) and m-FBXL4-R (TCAAGGATGCTGGACTTACCA), m-FBXL6-F (CCTGAGGGGTACCCGAGTTA) and m-FBXL6-R (GACACCACTGGACTTCCTCC), m-RPLP0-F (GGCGACCTGGAAGTCCAACT) and m-RPLP0-R (CCATCAGCACCACGGCCTTC), m-GUSB-F (TCATCTGGAATTTCGCCG) and m-GUSB-R (CTCGGCCCTGAACCGTGACCTC). Relative quantification was done by calculating the 2ΔΔCt value.

#### Plasmid construction

Plasmids used in this study are listed in the KEY RESOURCES TABLE. DNA cloning was performed either by DNA ligation using T4 DNA ligase (NEB), by recombination using In-Fusion HD cloning kit (Takara) or by site-directed mutagenesis using QuikChange II XL Site-Directed Mutagenesis Kit (Agilent Technologies).

Flag tag was fused in frame of the FBXL6 full length in order to have an expression of the tag at the C-terminal of the protein. C497R mutated FBXL6-Flag was obtained site-directed mutagenesis. MlsFBXL4-GFP and mlsFBXL6-GFP plasmids were created by PCR amplification of each mitochondrial leading sequence (mls) predicted by MITOPROT software https://ihg.gsf.de/ihg/mitoprot.html and subcloning into the AscI and XhoI sites of pCMV6-AC-GFP vector (Origene). Human FBXL6 was amplified by PCR and cloned into pCMV6-Entry from Origene (CAT#: PS100001). Cyto-GFP plasmid was obtained by subcloning of eGFP into BamHI and XhoI sites of pcDNA3 vector. Cyto-GFP-CAT plasmid was created by fusion PCR of eGFP and a C-terminal Ala-Thr tail (CAT tail): briefly a DNA fragment encoding eGFP fused at the C terminus with 10 Ala-Thr repeats followed by a stop codon, was amplified by PCR using primers eGFP-infusion-BamHI-F 5’-taccgagctcggatccgATG-3’ and eGFP-Fusion_CAT-tail-R 5’-gctgtcgctgtagcggttgcTGTAGCCGTTGCAGTGGCCTTGTACAGCTCGTCCATG-3’ and primers eGFP-Fusion_CAT-tail-F 5’-gcaaccgctacagcgacagcAACTGCCACTGCTACAGCGACTTAAatctcgagtctagaggg-3’ and eGFP-infusion-XhoI-R 5’-gccctctagactcgagATTTAA-3’ and cloned into BamHI and XhoI sites of pcDNA3. Mito-GFP and mito-GFP-CAT plasmids were created by PCR amplification of the mls of COX VIIIa and subcloning into cyto-GFP and cyto-GFP-CAT plasmid respectively. A DNA fragment encoding human MRPL45 was amplified from fibroblast cDNA by RT-PCR using primers MRPL45-BamHI-F 5’-CTTGGTACCGAGCTCGGATCCATGGCAGCCCCCATACCTCAAGG-3’ and MRPL45-XbaI-HA-R 5’-GGTTTAAACGGGCCCTCTAGATTAAGCGTAATCTGGAACATCGTATGGGTACTCGAGGGCTAGCTGAGGCTTC TGGG-3’ and cloned into BamHI and XbaI sites of pcDNA3 vector. MRPL45-CAT HA plasmid was obtained by PCR amplification of CAT tail and subcloning between MRPL45 DNA and HA-tag DNA into XhoI site of MRPL45-HA plasmid. MRPL45-HA-NS plasmid was obtained by site-directed mutagenesis of three stop codons after HA-tag using two sets of primers: primers NS-1-F 5’-tgttccagattacgctttatctagagggcccgtt-3’ and NS-1-R 5’-aacgggccctctagataaagcgtaatctggaaca-3’ and primers NS-2-F 5’-agagggcccgttttaacccgcttatcagcctcgac-3’ and NS-2-R 5’-gtcgaggctgataagcgggttaaaacgggccctct-3’. All plasmids were verified by sequencing before use.

#### Cell fractionation and trypsin accessibility assay

The different steps of cell fractionation were performed at 4°C. Cells were harvested in mitochondrial-isolation buffer [10 mM Tris HCl (pH 7.4), 210 mM mannitol, 70 mM sucrose, and 1 mM EDTA] supplemented with protease inhibitor (Sigma-Aldrich) and homogenized by passing through a 26-gauge syringe (20 strokes). The samples were centrifuged at 500g for 5 min at 4°C. Pellet were discarded. Supernatants were collected and centrifugated at 10,000g for 15 min at 4°C. The resulting supernatant was collected as the post mitochondrial supernatant fraction (PMS). The pellets were resuspended in 1 ml of isolation buffer and submitted to another round of centrifugation. The final pellet collected and solubilized in isolation buffer. An aliquote was saved as the mitochondria-enriched fraction/crude mitochondria (CM) and the remaining CM was treated with digitonin (0.5% final) for 15 min to remove all associated membranes. This fraction was centrifugated at 10,000g for 15 min at 4°C. Pellet was collected as the purified mitochondria (PM).

Trypsin accessibility assays were performed using enriched mitochondrial fractions. These fractions were exempt of protease inhibitors. Fractions were incubated with trypsin (0.01% final) for 0, 1, 5, 15 min and 15min with 0.5% of digitonin in presence of and reactions were stop with protease inhibitors cocktails and immediately centrifuged at 10000g for 10min. We collected the pellets that we resuspended in RIPA and sample buffer.

#### Immunoblots

Total-cell protein lysates and subcellular fractions were prepared using 2X sample buffer supplemented with protease inhibitors. Samples were analyzed by western blot using conventional methods. Briefly, 10 μg to 40 μg proteins were separated on 4-20% acrylamide gels by electrophoresis (120 V for 1 h). Proteins were transferred to PVDF membranes using TransBlot Turbo (BioRad) 10min at 2V. Membranes were blocked with 5% milk in phosphate-buffered saline (PBS)-Tween (0.05%) for 30 to 60 min. Proteins were detected using specific antibodies diluted in 5% milk/PBS-Tween (0.05%) and incubate from 1h to overnight according to the primary antibody used. As secondary antibodies we used horseradish peroxidase (HRP)-conjugated anti-rabbit or anti-mouse antibody (BioRad). HRP signals were visualized using chemiluminescent substrates (Thermo Fisher Scientific, Waltham, MA, USA) and acquired using a Chemidoc MP imaging system (BioRad). We also used IRDye 800CW Donkey anti-Rabbit or IRDye 680RD donkey anti-mouse IgG and revelation were performed using odyssey imaging system (Licor).

Images analyses were quantified using imageJ (NIH) for chemiluminescent signal or using Image Studio lite 5.2 (licor) for fluorescence signals.

#### BN-PAGE

Aggregation of endogenous MRPL45 was investigated in whole FBXL4-KO, FBXL6-KO, and control cell extracts using BN-PAGE. Samples were prepared at 4°C. Cells grown in 10 cm dishes were scrapped on ice and washed with cold PBS. Cell pellets were solubilized for 30 min on ice in 1X NativePAGE Sample Buffer (Invitrogen) containing protease inhibitors and 1.0% digitonine and then, centrifuged at 16,000x*g* for 20 min at 4 °C. Supernatants were collected and the protein concentration was determined using the BCA method. Prior to electrophoresis, the samples were supplemented with 0.25% Coomassie blue G (final concentration). For the first native dimension, 40 μg of protein was separated on 4–16% gradient native-PAGE gels (NativePAGE™ Novex^®^ Bis-Tris Gel from Invitrogen). Electrophoresis was performed using a light cathode buffer (NativePAGE 1X Running Buffer plus 0.02% G-250) and an anode buffer (NativePAGE 1X Running Buffer). Electrophoresis was performed at 150 v for about 2 h at room temperature.

Then, the gel strips were excised from the gels and incubated with 2X sample buffer containing 50 mM dithiothreitol (DTT) for 20 min. The lanes were placed on IPG tris/SDS gels (Biorad) and electrophoresis under denaturating conditions was performed using an SDS running buffer. Next, immunoblotting was performed following classical procedures.

#### Immunoprecipitation

For co-immunoprecipitation assays, HEK cells were transfected 48h before the assays. Cells transfected with β-galactosidase were used as control. Cells were scrapped and washed with cold PBS. Cell pellets were solubilized using IP lysis buffer (1% Triton X-100, 50 mM Tris (pH 7.4), 150 mM NaCl, 10 mM EDTA) supplemented with protease inhibitors. Solubilizations were performed for 20 min at 4°C with rotating mixing. Then, debris were removed by centrifugation for 20 min at 16,000g. Supernatants were incubated for 4h with anti-Flag or Ha agarose beads (Thermo Fisher Scientific and Merck) at 4°C with rotating mixing. Agarose beads were collected by centrifugation (30 seconds at 3000rpm). Beads were washed five times with cold PBS 0.05% Tween. Elution of proteins were performed using 2X Laemmli sample buffer (Merck) at 90°C for 5 min. Samples were analyzed by mass spectrometry as described above or immunoblots. For mass spectrometry, interactions were considered significant when each biological replicate displayed at least 2-fold enrichment as compared to control.

#### Proteome analyses by mass spectrometry

Ten μg of proteins were loaded on a 10% acrylamide SDS-PAGE gel and proteins were visualized by Colloidal Blue staining. Migration was stopped when samples had just entered the resolving gel and the unresolved region of the gel was cut into only one segment. Each SDS-PAGE band was cut into 1 mm x 1 mm gel pieces. Gel pieces were destained in 25 mM ammonium bicarbonate (NH4HCO3), 50% Acetonitrile (ACN) and shrunk in ACN for 10 min. After ACN removal, gel pieces were dried at room temperature. Proteins were first reduced in 10 mM dithiothreitol, 100 mM NH4HCO3 for 60 min at 56°C then alkylated in 100 mM iodoacetamide, 100 mM NH4HCO3 for 60 min at room temperature and shrunken in ACN for 10 min. After ACN removal, gel pieces were rehydrated with 50 mM NH4HCO3 for 10 min at room temperature. Before protein digestion, gel pieces were shrunken in ACN for 10 min and dried at room temperature. Proteins were digested by incubating each gel slice with 10 ng/μl of trypsin (V5111, Promega) in 40 mM NH4HCO3, rehydrated at 4°C for 10 min, and finally incubated overnight at 37°C. The resulting peptides were extracted from the gel by three steps: a first incubation in 40 mM NH4HCO3 for 15 min at room temperature and two incubations in 47.5% ACN, 5% formic acid for 15 min at room temperature. The three collected extractions were pooled with the initial digestion supernatant, dried in a SpeedVac, and resuspended with 0.1% formic acid for a final concentration of 0.05 μg/μL. NanoLC-MS/MS analysis were performed using an Ultimate 3000 RSLC Nano-UPHLC system (Thermo Scientific, USA) coupled to a nanospray Orbitrap Fusion™ Lumos Tribrid™ Mass Spectrometer (Thermo Fisher Scientific, California, USA). Each peptide extracts were loaded on a 300 μm ID x 5 mm PepMap C18 precolumn (Thermo Scientific, USA) at a flow rate of 10 μL/min. After a 3 min desalting step, peptides were separated on a 50 cm EasySpray column (75 μm ID, 2 μm C18 beads, 100 Å pore size, ES803A rev.2, Thermo Fisher Scientific) with a 4-40% linear gradient of solvent B (0.1% formic acid in 80% ACN) in 57 min. The separation flow rate was set at 300 nL/min. The mass spectrometer operated in positive ion mode at a 2.0 kV needle voltage. Data was acquired using Xcalibur 4.1 software in a data-dependent mode. MS scans (m/z 375-1500) were recorded at a resolution of R = 120000 (@ m/z 200) and an AGC target of 4×105 ions collected within 50 ms, followed by a top speed duty cycle of up to 3 seconds for MS/MS acquisition. Precursor ions (2 to 7 charge states) were isolated in the quadrupole with a mass window of 1.6 Th and fragmented with HCD@30% normalized collision energy. MS/MS data was acquired in the ion trap with rapid scan mode, AGC target of 3×103 ions and a maximum injection time of 35 ms. Selected precursors were excluded for 60 seconds. Protein identification and Label-Free Quantification (LFQ) were done in Proteome Discoverer 2.4. MS Amanda 2.0, Sequest HT and Mascot 2.5 algorithms were used for protein identification in batch mode by searching against a Uniprot Homo sapiens database (75 093 entries, release May 10, 2020). Two missed enzyme cleavages were allowed for the trypsin. Mass tolerances in MS and MS/MS were set to 10 ppm and 0.6 Da. Oxidation (M), acetylation (K) and deamidation (N, Q) were searched as dynamic modifications and carbamidomethylation, (C) as static modification.

Raw LC-MS/MS data were imported in Proline Web (Bouyssié et al. 2020) for feature detection, alignment, and quantification. Proteins identification was accepted only with at least 2 specific peptides with a pretty rank=1 and with a protein FDR value less than 1.0% calculated using the “decoy” option in Mascot. Label-free quantification of MS1 level by extracted ion chromatograms (XIC) was carried out with parameters indicated previously (Henriet et al. 2017). For expression analysis, the normalization was carried out on median of ratios. An inference of missing values was applied with 5% of the background noise when analyzing whole FBXL6-KO and control cell proteome. For immunoprecipitation data, no normalization or inference was applied. A Wilcoxon-Mann-Whitney test was applied to test the significance of the variation in relative protein abundances between experimental conditions.

#### Pulsed SILAC assay

Silac media was prepared as described by Ong and Mann (Ong and Mann 2007). Light media contains DMEM supplemented with 10% dialyzed FBS, Pen/Strep 1X; L-lysine (146mg.L^−1^) f, L-arginine (84mg.L^−1^) and proline (20mg.L^−1^). In heavy media, L-lysine and L-Arginine were replaced by L-lysine ^13^C6 (180mg.L^−1^) and ^13^C6 L-arginine (88mg.L^−1^), respectively. HEK cells were grown in light media for 2 weeks before uses. Two days before changing the media, cells were transfected with FBXL6-flag or control plasmid expressing β-galactosidase. At t=0h, cells were shifted to heavy media for 3h. Then, we performed a flag IP as described above and we saved an input fraction. Both input and pulled down fraction were subject to mass spectrometry analyses (see above). For pulsed SILAC, ^13^C-K and ^13^C-R were searched as static modifications.

#### Turbo ID and mass spectrometry analysis

TurboID proximity labeling is based on previously described protocol (Cho et al., 2020). Briefly, TurboID was fused to the C-ternimus of GFP or FBXL6. HCT116 stable cell line expressing FBXL6-TurboID and TurboID-GFP were grown for 30 minutes and 16 hours in presence of 50 μM of biotin (Sigma Aldrich). About 20 million cells (two 10-cm plates) were first washed two times with ice-cold PBS and lysed in 1 mL of RIPA buffer (50 mM Tris HCL, pH 7.4; 150 mM NaCl, 0.1% SDS, 0.5% Sodium Deoxycholate, 1% triton 100X) for 10 minutes on ice. The extract is sonicated 3 times at 20 % amplitude for 20 seconds and cleared by centrifugation at 12 000 g for 10 minutes. The supernatant kept to pulldown biotinylated proteins by the addition of 100 μL of streptavidin-coupled magnetic beads slurry (Streptavidin Mag Sepharose, GE Healthcare) previously washed and equilibrated. Samples are put on rotating wheel overnight at 4°C. The beads are washed 5 minutes at RT on rotating wheel, twice with RIPA buffer, once with 1M KCL, with 0.1M Na2CO3 and with 2M urea in 10mM Tris-HCL pH8. Beads are washed again twice with RIPA buffer and with 50 mM NH4HCO3. A fraction of the beads (5%) is boiled in Laemmli buffer for western blotting and the remaining beads are analyzed by LC/MS-MS.

For mass-spectrometry analyses, magnetic beads were extensively washed in 50 mM ammonium bicarbonate and proteins were digested directly on the beads in 2 consecutive steps with sequencing-grade porcine trypsin (Promega, Fitchburg, MA, USA).Peptides generated after trypsin digestion were analysed by nanoLC-MS/MS on a QExactive + mass spectrometer coupled to an EASY-nanoLC-1000 (Thermo-Fisher Scientific, USA). Peptides were identified with Mascot algorithm (Matrix Science, London, UK): the data were searched against the Swissprot updated databases with Homo sapiens taxonomies using the software’s decoy strategy. Mascot 2.6 and Swissprot version 2020_05 (20386 sequences) were used for the experiment. Mascot identifications were imported into Proline 1.4 software (Bouyssié et al. 2020) where they were validated using the following settings: PSM score <=25, Mascot pretty rank < = 1, FDR < = 1% for PSM scores, FDR < = 1% and for protein set scores. The total number of MS/MS fragmentation spectra was used to quantify each protein.

For the statistical analysis of the data, we compared the different samples against the negative controls using R v3.5.2 and R-studio. The spectral counts from 3 replicates were normalized according to the DESeq2 normalization method (i.e., median of ratios method) and EdgeR was used to perform a negative-binomial test. For each identified protein, an adjusted p-value (adjp) corrected by Benjamini– Hochberg was calculated, as well as a protein fold-change (FC). The results are presented in a Volcano plot using protein log2 fold changes and their corresponding adjusted (-log10adjp) to highlight up regulated and down regulated proteins.

#### Proteomics data availability

The mass spectrometry proteomics data have been deposited to the ProteomeXchange Consortium via the PRIDE partner repository (Deutsch et al. 2020). These data are accessible through the following identifiers:

Proteomic analysis of FBXL6 Knockdown cells proteome

Project accession: PXD037252

Reviewer account details:

Username

Password: uke34Jh3

Proteomic analysis of FBXL6 interacting proteome

Project accession: PXD037248

Reviewer account details:

Username

Password: FIO5OMZb

Analyses of FBXL6 interaction with newly synthesized proteins.

Project accession: PXD037602

Reviewer account details:

Username

Password: ZD7EJ1rL

Analysis of biotinylated proteins in cells expressing GFP-TurboID

Project accession: PXD027122

Username

Password: kzY32FDg

Analysis of biotinylated proteins in cells expressing FBXL6-TurboID cells

Project accession: PXD028161

Username

Password: LN8C1uPJ

#### Immunofluorescence microscopy

Cells were grown on glass coverslips and transfected using Fugene HD (Promega). After 24 h-48h, cells were fixed with 4% paraformaldehyde for 15 min at room temperature (all of the following steps were done at room temperature) and then, PFA was washed out with phosphate-buffered saline. Cells were permeabilized in 0.2% Triton X-100–phosphate-buffered saline for 15 min and saturated with 10% bovine serum albumin in PBS for 45 min. Primary antibodies were incubated for 2 h in the blocking buffer (10% BSA in PBS). Three washes of 10 minutes with blocking buffer were performed before incubation with secondary antibodies for 45 min. Following three washes of 10 minutes with PBS, immunostainings were visualized with a Zeiss microscope. Images were acquired using a Zeiss microscope (AxioZision; Carl Zeiss, OberKOchen, Germany) with a 63× objective. Z-sections (interval: 0.2 μm) covering the entire depth of the cell were acquired. Quantifications (levels of fluorescence and Pearson’s correlation coefficient) were performed using Zeiss co-localization software (Carl Zeiss).

#### Oxygen consumption analyses and ATP measurement

To measured cellular oxygen consumption rate, cells were grown on 10cm dishes and 24h before testing, growing media were replaced by no glucose media supplemented with glutamine. Cells were collected and resuspended to 1 × 10^6^ cells/mL in glutamine media.

Cellular respiration was measured from 1 × 10^6^ cells/mL to 5 × 10^6^ cells/mL at 37°C. Mitochondrial oxygen-consumption assays were performed using the high-resolution respirometry system Oxygraph-2k (Oroboros). The basal respiration, the oligomycin- and the FCCP-dependent respiration were successively recorded over the time.

To assay ATP content, cells were seeded in a 96-well white plate with a clear bottom at 8000 cells per well. The following day, the intracellular ATP content was measured by using the bioluminescent ATP Kit CellTiter-Glo^®^ 2.0 (Promega) following the manufacturer’s instructions. The part of ATP produced by mitochondria was calculated as the difference between the total ATP content and the amount of ATP measured in the presence of the ATP synthase inhibitor oligomycin. Cells were incubated 30 min with 3 μM oligomycin to inhibit mitochondrial respiration and linked-ATP synthesis before adding the CellTiter-Glo^®^ 2.0 Reagent to the wells. Luminescence was measured by a multiplate reader (Luminoskan Microplate Luminometer, Thermo Fisher Scientific, Waltham, MA, USA). Standardization was performed with known quantities of standard ATP in the same conditions.

#### Cell Cycle assays

Cell was incubated for 48h in glucose or galactose medium. Edu labelling was made according manufacturer recommendation (Thermofisher Click it Edu). Briefly, Edu 10μM was added for 1.5 hours before pellet cell Fixed and permeabilized cell was incubated with Rnase for 2 hours 37°C before Alexa 647 Click it Edu Reaction. Then, Cell was incubated 20 min for DNA content with a 10% dilution IP/Rnase (FxCycle, Invitrogen). Before Facs analysis (BD Accuri C6 plus).

#### Quantification and statistical analysis

All values represent the mean ± SEM. Statistical analyses were performed using the Prism 9 software (GraphPad). Normality distribution was assessed using the D’Agostino-Pearson test. The p-value, tests, and post hoc tests are cited in the figure legend.

Gene Ontology analyses were performed using DAVID 6.8 (Huang, Sherman, and Lempicki 2009). We selected GOTERM_BP_DIRECT pathways containing at least three proteins. For Hela cells we used background from Robin et al (Robin et al. 2019) and for HEK cells Lavado-Garcia et al (Lavado-García et al. 2020). We used proteins presenting significantly (p<0.05) increased or decreased expression following mass spectrometry analyses.

## References

Alsina, David, Oleksandr Lytovchenko, Aleksandra Schab, Ilian Atanassov, Florian A Schober, Min Jiang, Camilla Koolmeister, et al. 2020. “FBXL 4 Deficiency Increases Mitochondrial Removal by Autophagy EMBO Molecular Medicine 12 (7): 1–16. https://doi.org/10.15252/emmm.201911659.

Ansar, Muhammad, Sohail Aziz Paracha, Alessandro Serretti, Muhammad T. Sarwar, Jamshed Khan, Emmanuelle Ranza, Emilie Falconnet, et al. 2019. “Biallelic Variants in FBXL3 Cause Intellectual Disability, Delayed Motor Development and Short Stature.” Human Molecular Genetics 28 (6): 972–79. https://doi.org/10.1093/hmg/ddy406.

Bengtson, Mario H., and Claudio A.P. Joazeiro. 2010. “Role of a Ribosome-Associated E3 Ubiquitin Ligase in Protein Quality Control.” Nature 467 (7314): 470–73. https://doi.org/10.1038/nature09371.

Bonnen, Penelope E., John W. Yarham, Arnaud Besse, Ping Wu, Eissa A. Faqeih, Ali Mohammad Al-Asmari, Mohammad A.M. Saleh, et al. 2013. “Mutations in FBXL4 Cause Mitochondrial Encephalopathy and a Disorder of Mitochondrial DNA Maintenance.” American Journal of Human Genetics 93 (3): 471–81. https://doi.org/10.1016/j.ajhg.2013.07.017.

Boos, Felix, Lena Krämer, Carina Groh, Ferris Jung, Per Haberkant, Frank Stein, Florian Wollweber, et al. 2019. “Mitochondrial Protein-Induced Stress Triggers a Global Adaptive Transcriptional Programme.” Nature Cell Biology 21 (4): 442–51. https://doi.org/10.1038/s41556-019-0294-5.

Bouyssié, David, Anne Marie Hesse, Emmanuelle Mouton-Barbosa, Magali Rompais, Charlotte MacRon, Christine Carapito, Anne Gonzalez De Peredo, et al. 2020. “Proline: An Efficient and User-Friendly Software Suite for Large-Scale Proteomics.” Bioinformatics 36 (10): 3148–55. https://doi.org/10.1093/bioinformatics/btaa118.

Branon, Tess C., Justin A. Bosch, Ariana D. Sanchez, Namrata D. Udeshi, Tanya Svinkina, Steven A. Carr, Jessica L. Feldman, Norbert Perrimon, and Alice Y. Ting. 2018. “Efficient Proximity Labeling in Living Cells and Organisms with TurboID.” Nature Biotechnology 36 (9): 880–98. https://doi.org/10.1038/nbt.4201.

Chu, Jessie, Nancy A. Hong, Claudio A. Masuda, Brian V. Jenkins, Keats A. Nelms, Christopher C. Goodnow, Richard J. Glynne, et al. 2009. “A Mouse Forward Genetics Screen Identifies LISTERIN as an E3 Ubiquitin Ligase Involved in Neurodegeneration.” Proceedings of the National Academy of Sciences of the United States of America 106 (7): 2097–2103. https://doi.org/10.1073/pnas.0812819106.

Defenouillère, Quentin, Elodie Zhang, Abdelkader Namane, John Mouaikel, Alain Jacquier, and Micheline Fromont-Racine. 2016. “Rqcl and Ltn1 Prevent C-Terminal Alanine-Threonine Tail (Cat-Tail)-Induced Protein Aggregation by Efficient Recruitment of Cdc48 on Stalled 60s Subunits.” Journal of Biological Chemistry 291 (23): 12245–53. https://doi.org/10.1074/jbc.M116.722264.

Deutsch, Eric W., Nuno Bandeira, Vagisha Sharma, Yasset Perez-Riverol, Jeremy J. Carver, Deepti J. Kundu, David García-Seisdedos, et al. 2020. “The ProteomeXchange Consortium in 2020: Enabling 舖big Data舗 Approaches in Proteomics.” Nucleic Acids Research 48 (D1): D1145–52. https://doi.org/10.1093/nar/gkz984.

Duttler, Stefanie, Sebastian Pechmann, and Judith Frydman. 2013. “Principles of Cotranslational Ubiquitination and Quality Control at the Ribosome.” Molecular Cell 50 (3): 379–93. https://doi.org/10.1016/j.molcel.2013.03.010.

El-Hattab, Ayman W., Hongzheng Dai, Mohammed Almannai, Julia Wang, Eissa A. Faqeih, Ali Al Asmari, Mohammed A.M. Saleh, et al. 2017. “Molecular and Clinical Spectra of FBXL4 Deficiency.” Human Mutation 38 (12): 1649–59. https://doi.org/10.1002/humu.23341.

Finley, Daniel, Helle D. Ulrich, Thomas Sommer, and Peter Kaiser. 2012. “The Ubiquitin-Proteasome System of Saccharomyces Cerevisiae.” Genetics 192 (2): 319–60. https://doi.org/10.1534/genetics.112.140467.

Gai, Xiaowu, Daniele Ghezzi, Mark A. Johnson, Caroline A. Biagosch, Hanan E. Shamseldin, Tobias B. Haack, Aurelio Reyes, et al. 2013. “Mutations in FBXL4, Encoding a Mitochondrial Protein, Cause Early-Onset Mitochondrial Encephalomyopathy.” American Journal of Human Genetics 93 (3): 482–95. https://doi.org/10.1016/j.ajhg.2013.07.016.

Gold, Vicki AM, Piotr Chroscicki, Piotr Bragoszewski, and Agnieszka Chacinska. 2017. “Visualization of Cytosolic Ribosomes on the Surface of Mitochondria by Electron Cryo-tomography.” EMBO Reports 18 (10): 1786–1800. https://doi.org/10.15252/embr.201744261.

Haaften-Visser, Désirée Y. Van, Magdalena Harakalova, Enric Mocholi, Joris M. Van Montfrans, Abdul Elkadri, Ester Rieter, Karoline Fiedler, et al. 2017. “Ankyrin Repeat and Zinc-Finger Domain-Containing 1 Mutations Are Associated with Infantile-Onset Inflammatory Bowel Disease.” Journal of Biological Chemistry 292 (19): 7904–20. https://doi.org/10.1074/jbc.M116.772038.

Hartl, F. Ulrich, and Manajit Hayer-Hartl. 2002. “Protein Folding. Molecular Chaperones in the Cytosol: From Nascent Chain to Folded Protein.” Science 295 (5561): 1852–58. https://doi.org/10.1126/science.1068408.

Henriet, Elodie, Aya Abou Hammoud, Jean William Dupuy, Benjamin Dartigues, Zakaria Ezzoukry, Nathalie Dugot-Senant, Thierry Leste-Lasserre, et al. 2017. “Argininosuccinate Synthase 1 (ASS1): A Marker of Unclassified Hepatocellular Adenoma and High Bleeding Risk.” Hepatology. https://doi.org/10.1002/hep.29336.

Heo, Jin Mi, Nurit Livnat-Levanon, Eric B. Taylor, Kevin T. Jones, Noah Dephoure, Julia Ring, Jianxin Xie, et al. 2010. “A Stress-Responsive System for Mitochondrial Protein Degradation.” Molecular Cell 40 (3): 465–80. https://doi.org/10.1016/j.molcel.2010.10.021.

Hirano, Arisa, Kanae Yumimoto, Ryosuke Tsunematsu, Masaki Matsumoto, Masaaki Oyama, Hiroko Kozuka-Hata, Tomoki Nakagawa, Darin Lanjakornsiripan, Keiichi I. Nakayama, and Yoshitaka Fukada. 2013. “FBXL21 Regulates Oscillation of the Circadian Clock through Ubiquitination and Stabilization of Cryptochromes.” Cell 152 (5): 1106–18. https://doi.org/10.1016/j.cell.2013.01.054.

Huang, Da Wei, Brad T. Sherman, and Richard A. Lempicki. 2009. “Systematic and Integrative Analysis of Large Gene Lists Using DAVID Bioinformatics Resources.” Nature Protocols 4 (1): 44–57. https://doi.org/10.1038/nprot.2008.211.

Huemer, Martina, Daniela Karall, Anna Schossig, Jose E. Abdenur, Fatma Al Jasmi, Caroline Biagosch, Felix Distelmaier, et al. 2015. “Clinical, Morphological, Biochemical, Imaging and Outcome Parameters in 21 Individuals with Mitochondrial Maintenance Defect Related to FBXL4 Mutations.” Journal of Inherited Metabolic Disease 38 (5): 905–14. https://doi.org/10.1007/s10545-015-9836-6.

Izawa, Toshiaki, Sae Hun Park, Liang Zhao, F. Ulrich Hartl, and Walter Neupert. 2017. “Cytosolic Protein Vms1 Links Ribosome Quality Control to Mitochondrial and Cellular Homeostasis.” Cell 171 (4): 890–903.e18. https://doi.org/10.1016/j.cell.2017.10.002.

Jaiswal, Himjyot, Charlotte Conz, Hendrik Otto, Tina Wölfle, Edith Fitzke, Matthias P. Mayer, and Sabine Rospert. 2011. “The Chaperone Network Connected to Human Ribosome-Associated Complex.” Molecular and Cellular Biology 31 (6): 1160–73. https://doi.org/10.1128/mcb.00986-10.

Joazeiro, Claudio A.P. 2019. “Mechanisms and Functions of Ribosome-Associated Protein Quality Control.” Nature Reviews Molecular Cell Biology 20 (6): 368–83. https://doi.org/10.1038/s41580-019-0118-2.

Karbowski, Mariusz, Albert Neutzner, and Richard J. Youle. 2007. “The Mitochondrial E3 Ubiquitin Ligase MARCH5 Is Required for Drp1 Dependent Mitochondrial Division.” Journal of Cell Biology 178 (1): 71–84. https://doi.org/10.1083/jcb.200611064.

Kellems, Rod E., and Ronald A. Butow. 1974. “Cytoplasmic Type 80 S Ribosomes Associated with Yeast Mitochondria.” Journal of Biological Chemistry. https://doi.org/10.1016/s0021-9258(19)42673-2.

Kostova, Kamena K., Kelsey L. Hickey, Beatriz A. Osuna, Jeffrey A. Hussmann, Adam Frost, David E. Weinberg, and Jonathan S. Weissman. 2017. “CAT-Tailing as a Fail-Safe Mechanism for Efficient Degradation of Stalled Nascent Polypeptides.” Science 357 (6349): 414–17. https://doi.org/10.1126/science.aam7787.

Kuroha, Kazushige, Alexandra Zinoviev, Christopher U.T. Hellen, and Tatyana V. Pestova. 2018. “Release of Ubiquitinated and Non-Ubiquitinated Nascent Chains from Stalled Mammalian Ribosomal Complexes by ANKZF1 and Ptrh1.” Molecular Cell 72 (2): 286–302.e8. https://doi.org/10.1016/j.molcel.2018.08.022.

Lavado-García, Jesús, Inmaculada Jorge, Laura Cervera, Jesús Vázquez, and Francesc Gòdia. 2020. “Multiplexed Quantitative Proteomic Analysis of HEK293 Provides Insights into Molecular Changes Associated with the Cell Density Effect, Transient Transfection, and Virus-Like Particle Production.” Journal of Proteome Research 19 (3): 1085–99. https://doi.org/10.1021/acs.jproteome.9b00601.

Lavie, Julie, Harmony De Belvalet, Sessinou Sonon, Ana Madalina Ion, Elodie Dumon, Su Melser, Didier Lacombe, Jean William Dupuy, Claude Lalou, and Giovanni Bénard. 2018. “Ubiquitin-Dependent Degradation of Mitochondrial Proteins Regulates Energy Metabolism.” Cell Reports 23 (10): 2852–63. https://doi.org/10.1016/j.celrep.2018.05.013.

Lazarou, Michael, Seok Min Jin, Lesley A. Kane, and Richard J. Youle. 2012. “Role of PINK1 Binding to the TOM Complex and Alternate Intracellular Membranes in Recruitment and Activation of the E3 Ligase Parkin.” Developmental Cell 22 (2): 320–33. https://doi.org/10.1016/j.devcel.2011.12.014.

Lesnik, Chen, Yifat Cohen, Avigail Atir-Lande, Maya Schuldiner, and Yoav Arava. 2014. “OM14 Is a Mitochondrial Receptor for Cytosolic Ribosomes That Supports Co-Translational Import into Mitochondria.” Nature Communications 5: 1–10. https://doi.org/10.1038/ncomms6711.

Li, Wei, Mario H. Bengtson, Axel Ulbrich, Akio Matsuda, Venkateshwar A. Reddy, Anthony Orth, Sumit K. Chanda, Serge Batalov, and Claudio A.P. Joazeiro. 2008. “Genome-Wide and Functional Annotation of Human E3 Ubiquitin Ligases Identifies MULAN, a Mitochondrial E3 That Regulates the Organelle舗s Dynamics and Signaling.” PLoS ONE 3 (1). https://doi.org/10.1371/journal.pone.0001487.

Mårtensson, Christoph U., Chantal Priesnitz, Jiyao Song, Lars Ellenrieder, Kim Nguyen Doan, Felix Boos, Alessia Floerchinger, et al. 2019. “Mitochondrial Protein Translocation-Associated Degradation.” Nature 569 (7758): 679–83. https://doi.org/10.1038/s41586-019-1227-y.

Mason, Bethany, and Heike Laman. 2020. “The FBXL Family of F-Box Proteins: Variations on a Theme: The FBXL Family of F-Box Proteins.” Open Biology 10 (11). https://doi.org/10.1098/rsob.200319.

Melser, Su, Etienne Hébert Chatelain, Julie Lavie, Walid Mahfouf, Caroline Jose, Emilie Obre, Susan Goorden, et al. 2013. “Rheb Regulates Mitophagy Induced by Mitochondrial Energetic Status.” Cell Metabolism 17 (5): 719–30. https://doi.org/10.1016/j.cmet.2013.03.014.

Mohanraj, Karthik, Michal Wasilewski, Cristiane Benincá, Dominik Cysewski, Jaroslaw Poznanski, Paulina Sakowska, Zaneta Bugajska, et al. 2019. “Inhibition of Proteasome Rescues a Pathogenic Variant of Respiratory Chain Assembly Factor COA7.” EMBO Molecular Medicine 11 (5): 1–21. https://doi.org/10.15252/emmm.201809561.

Ong, Shao En, and Matthias Mann. 2007. “A Practical Recipe for Stable Isotope Labeling by Amino Acids in Cell Culture (SILAC).” Nature Protocols 1 (6): 2650–60. https://doi.org/10.1038/nprot.2006.427.

Pechmann, Sebastian, Felix Willmund, and Judith Frydman. 2013. “The Ribosome as a Hub for Protein Quality Control.” Molecular Cell 49 (3): 411–21. https://doi.org/10.1016/j.molcel.2013.01.020.

Phu, Lilian, Christopher M. Rose, Joy S. Tea, Christopher E. Wall, Erik Verschueren, Tommy K. Cheung, Donald S. Kirkpatrick, and Baris Bingol. 2020. “Dynamic Regulation of Mitochondrial Import by the Ubiquitin System.” Molecular Cell 77 (5): 1107–1123.e10. https://doi.org/10.1016/j.molcel.2020.02.012.

Pisareva, Vera P., Maxim A. Skabkin, Christopher U.T. Hellen, Tatyana V. Pestova, and Andrey V. Pisarev. 2011. “Dissociation by Pelota, Hbs1 and ABCE1 of Mammalian Vacant 80S Ribosomes and Stalled Elongation Complexes.” EMBO Journal 30 (9): 1804–17. https://doi.org/10.1038/emboj.2011.93.

Rath, Sneha, Rohit Sharma, Rahul Gupta, Tslil Ast, Connie Chan, Timothy J. Durham, Russell P. Goodman, et al. 2021. “MitoCarta3.0: An Updated Mitochondrial Proteome Now with Sub-Organelle Localization and Pathway Annotations.” Nucleic Acids Research 49 (D1): D1541–47. https://doi.org/10.1093/nar/gkaa1011.

Reitzer, L. J., B. M. Wice, and D. Kennell. 1979. “Evidence That Glutamine, Not Sugar, Is the Major Energy Source for Cultured HeLa Cells.” Journal of Biological Chemistry 254 (8): 2669–76. https://doi.org/10.1016/s0021-9258(17)30124-2.

Robin, Thibault, Amos Bairoch, Markus Müller, Frédérique Lisacek, and Lydie Lane. 2019. “Correction to: Large-Scale Reanalysis of Publicly Available Hela Cell Proteomics Data in the Context of the Human Proteome Project (Journal of Proteome Research (2018) 17:12 (4160-4170) DOI: 10.1021/Acs.Jproteome.8b00392).” Journal of Proteome Research 18 (4): 1926–27. https://doi.org/10.1021/acs.jproteome.9b00113.

Sabouny, Rasha, Rachel Wong, Laurie Lee-Glover, Steven C. Greenway, David S. Sinasac, Aneal Khan, and Timothy E. Shutt. 2019. “Characterization of the C584R Variant in the MtDNA Depletion Syndrome Gene FBXL4, Reveals a Novel Role for FBXL4 as a Regulator of Mitochondrial Fusion.” Biochimica et Biophysica Acta-Molecular Basis of Disease 1865 (11): 165536. https://doi.org/10.1016/j.bbadis.2019.165536.

Shao, Sichen, Alan Brown, Balaji Santhanam, and Ramanujan S. Hegde. 2015. “Structure and Assembly Pathway of the Ribosome Quality Control Complex.” Molecular Cell 57 (3): 433–44. https://doi.org/10.1016/j.molcel.2014.12.015.

Shen, Peter S, Joseph Park, Yidan Qin, Xueming Li, Krishna Parsawar, Matthew H Larson, James Cox, et al. 2015. “Elongation of Nascent Chains.” Science (New York, N.Y.) 1 (1): 1–2.

Smith, Anthony C., and Alan J. Robinson. 2016. “MitoMiner v3.1, an Update on the Mitochondrial Proteomics Database.” Nucleic Acids Research 44 (D1): D1258–61. https://doi.org/10.1093/nar/gkv1001.

Thrun, Anna, Aitor Garzia, Yu Kigoshi-Tansho, Pratik R. Patil, Charles S. Umbaugh, Teresa Dallinger, Jia Liu, et al. 2021. “Convergence of Mammalian RQC and C-End Rule Proteolytic Pathways via Alanine Tailing.” Molecular Cell 81 (10): 2112–2122.e7. https://doi.org/10.1016/j.molcel.2021.03.004.

Tian, Guiyou, Cheng Hu, Yun Yun, Wensheng Yang, Wolfgang Dubiel, Yabin Cheng, and Dieter A Wolf. 2021. “Dual Roles of HSP70 Chaperone HSPA1 in Quality Control of Nascent and Newly Synthesized Proteins.” The EMBO Journal, 1–23. https://doi.org/10.15252/embj.2020106183.

Verma, Rati, Kurt M. Reichermeier, A. Maxwell Burroughs, Robert S. Oania, Justin M. Reitsma, L. Aravind, and Raymond J. Deshaies. 2018. “Vms1 and ANKZF1 Peptidyl-TRNA Hydrolases Release Nascent Chains from Stalled Ribosomes.” Nature 557 (7705): 446–51. https://doi.org/10.1038/s41586-018-0022-5.

Wrobel, Lidia, Ulrike Topf, Piotr Bragoszewski, Sebastian Wiese, Malgorzata E. Sztolsztener, Silke Oeljeklaus, Aksana Varabyova, et al. 2015. “Mistargeted Mitochondrial Proteins Activate a Proteostatic Response in the Cytosol.” Nature 524 (7566): 485–88. https://doi.org/10.1038/nature14951.

Wu, Zhihao, Yan Wang, Junghyun Lim, Boxiang Liu, Yanping Li, Rasika Vartak, Trisha Stankiewicz, Stephen Montgomery, and Bingwei Lu. 2018. “Ubiquitination of ABCE1 by NOT4 in Response to Mitochondrial Damage Links Co-Translational Quality Control to PINK1-Directed Mitophagy.” Cell Metabolism 28 (1): 130–144.e7. https://doi.org/10.1016/j.cmet.2018.05.007.

Young, Jason C, Nicholas J Hoogenraad, and F Ulrich Hartl. 2005. “Contents, Ed. Board + Forthc. Articles.” Trends in Biochemical Sciences 30 (3): i. https://doi.org/10.1016/s0968-0004(05)00043-5.

