## supplemental figures for "The E3 ubiquitin ligase FBXL6 controls the quality of newly synthesized mitochondrial ribosomal proteins"

**Table S1** : Excel proteomic FBXL6 KO vs ctrl

**Table S2 :** Excel proteomic IP ctrl vs FBXL6

**Table S3 :** Excel proteomic IP ctrl vs FBXL6 GO analyses

**Table S4 :** Excel proteomic turboID ctrl vs FBXL6

**Figure S1**

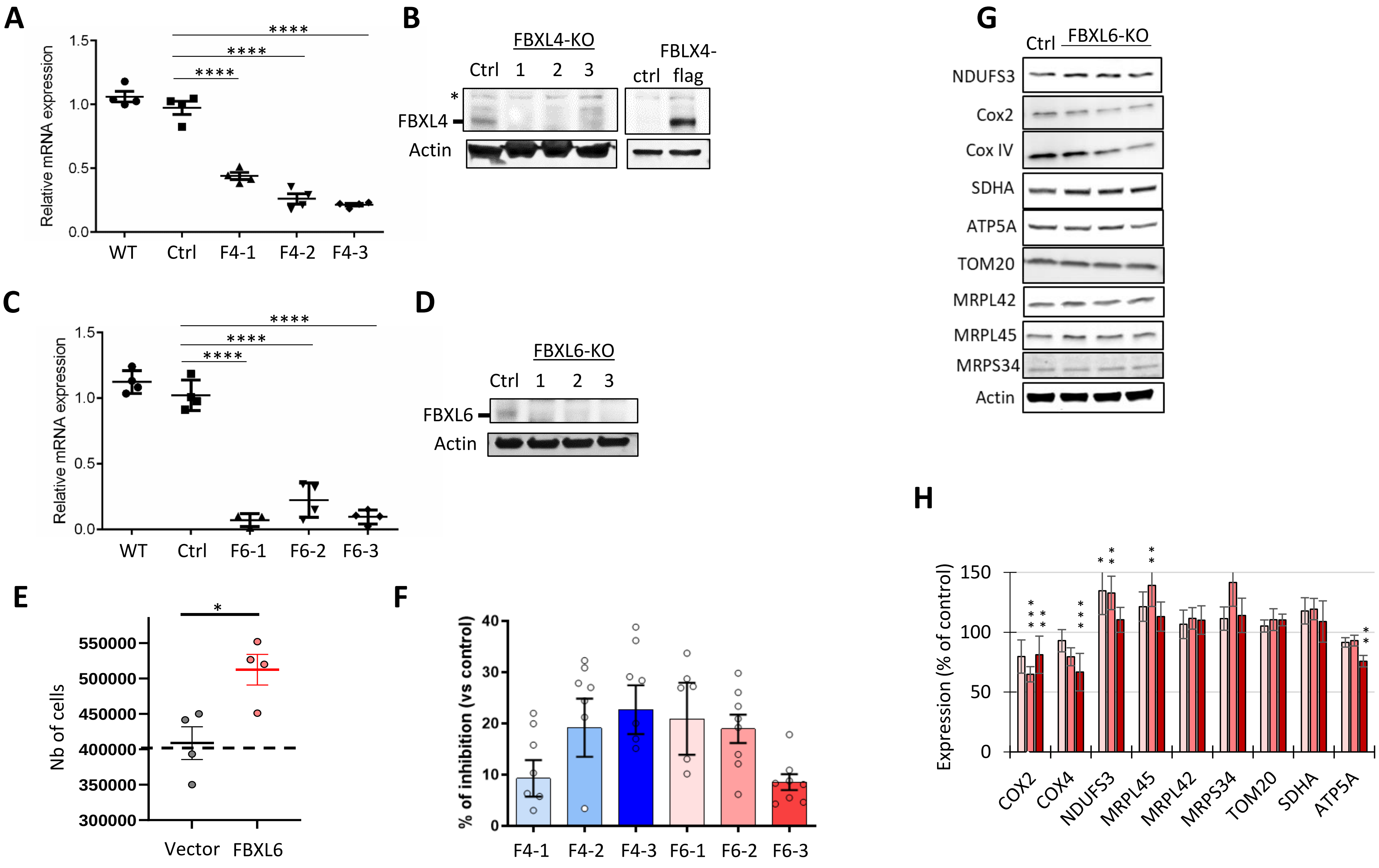

A
Relative mRNA expression
1.5
1.0
0.5
0.0
****
WT
Ctrl
F4-1
F4-2
F4-3
B
FBXL4-KO
Ctrl 1 2 3
FBLX4-
ctrl flag
*
FBXL4
Actin
C
Relative mRNA expression
WT
Ctrl
F6-1
F6-2
F6-3
D
FBXL6-KO
Ctrl 1 2 3
FBXL6
Actin
E
Nb of cells
550000
500000
450000
400000
350000
300000
Vector
FBXL6
F
% of inhibition (vs control)
40
30
20
10
0
F4-1
F4-2
F4-3
F6-1
F6-2
F6-3
G
Ctrl
FBXL6-KO
NDUFS3
Cox2
Cox IV
SDHA
ATP5A
TOM20
MRPL42
MRPL45
MRPS34
Actin
H
Expression (% of control)
150
100
50
0
COX2
COX4
NDUFS3
MRPL45
MRPL42
MRPS34
TOM20
SDHA
ATP5A

**Figure S2**

| Gene Symbol | Mito Evidence Mass-Spec Studies | Mito Evidence GFP | Mito Evidence Mass-Spec Experiments | Mito Evidence GO Annotation | Mito Evidence IMPI | Mito Evidence MitoCarta | Mito Evidence IMPI score | Mito Evidence Human Protein Atlas | Mito Targeting Seq iPSORT | Mito Targeting Seq MitoProt | Mito Targeting Seq TargetP |
|---|---|---|---|---|---|---|---|---|---|---|---|
| FBXL6 | 0 | 0 | 0 | false | Predicted NOT mitochondrial | false | 0.627204893 | false | 1.0 | 0.9725 | 0.624 |
| FBXL4 | 0 | 0 | 0 | true | Known mitochondrial | true | 0.791611975 | false | 0.0 | 0.9427 | 0.535 |

Figure S3

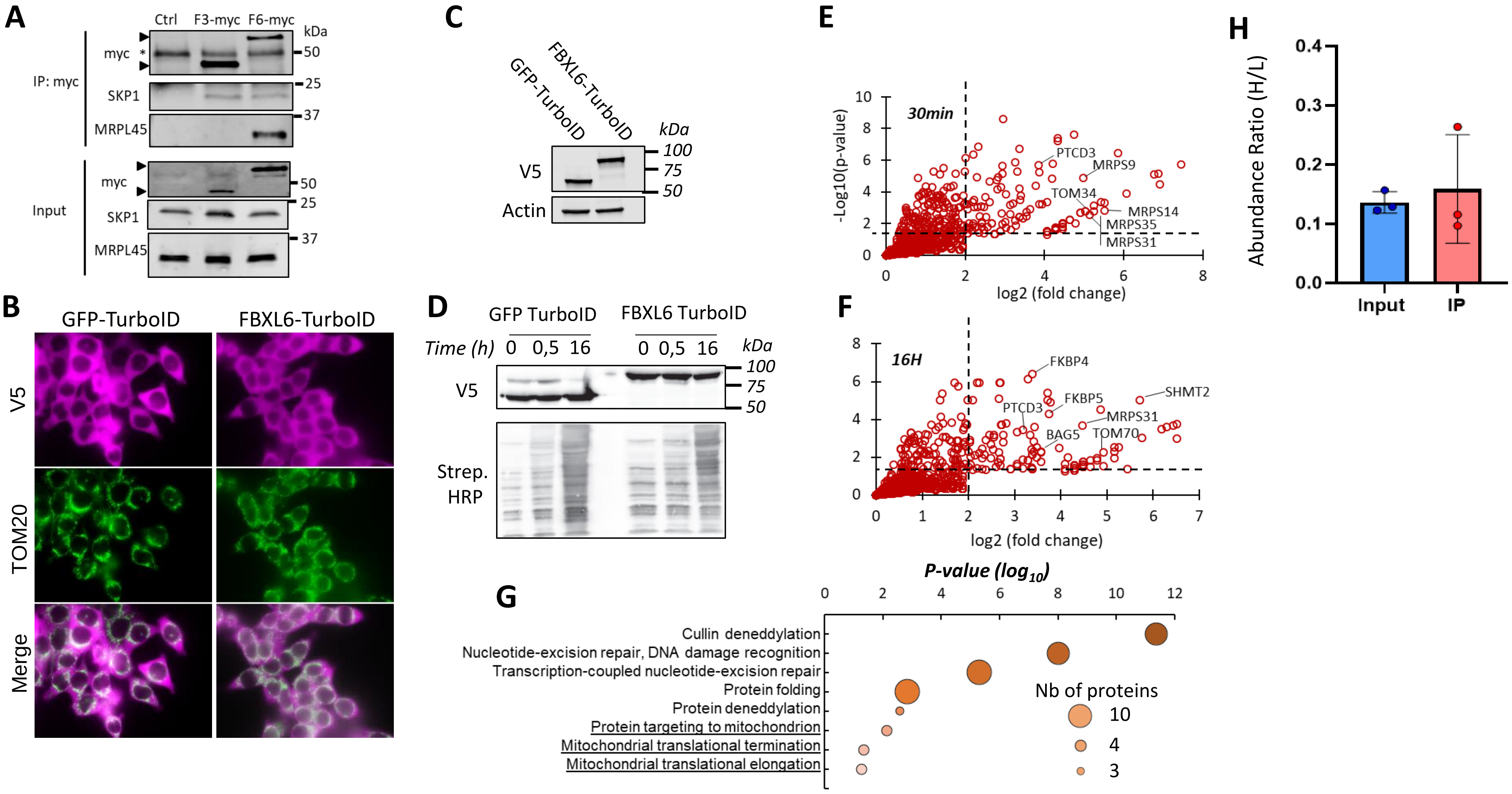
A
Ctrl
F3-myc
F6-myc
kDa
IP: myc
myc
SKP1
MRPL45
Input
myc
SKP1
MRPL45
50
25
37
B
GFP-TurboID
FBXL6-TurboID
V5
TOM20
Merge
C
GFP-TurboID
FBXL6-TurboID
kDa
V5
Actin
100
75
50
D
GFP TurboID
FBXL6 TurboID
Time (h)
0 0,5 16
0 0,5 16
kDa
V5
Strep. HRP
100
75
50
E
30min
-Log10(p-value)
log2 (fold change)
PTCD3
MRPS9
TOM34
MRPS14
MRPS35
MRPS31
F
16H
log2 (fold change)
FKBP4
SHMT2
FKBP5
PTCD3
MRPS31
BAG5
TOM70
G
P-value (log10)
Cullin deneddylation
Nucleotide-excision repair, DNA damage recognition
Transcription-coupled nucleotide-excision repair
Protein folding
Protein deneddylation
Protein targeting to mitochondrion
Mitochondrial translational termination
Mitochondrial translational elongation
Nb of proteins
10
4
3
H
Abundance Ratio (H/L)
Input
IP

#### Figure S4

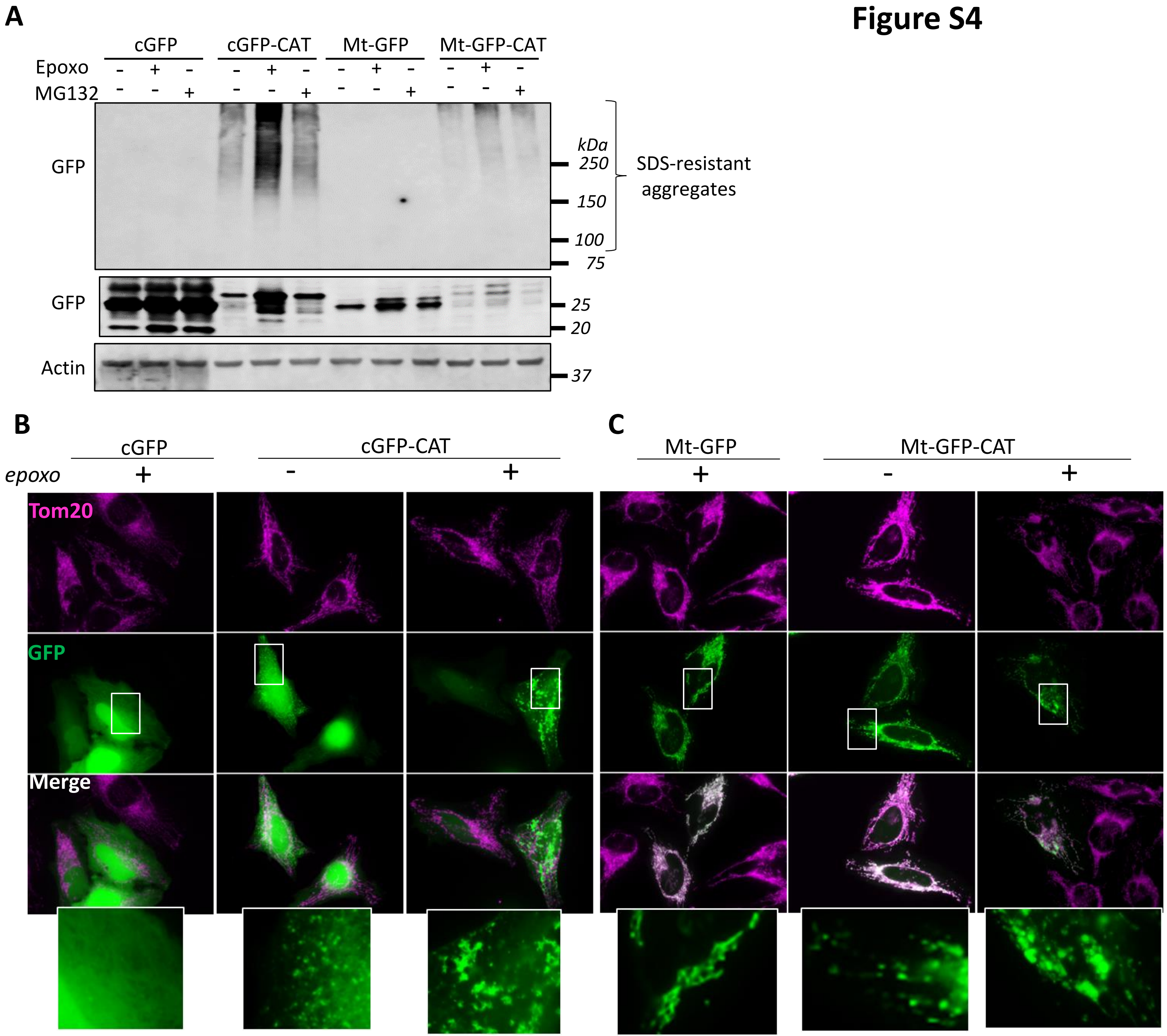
A
cGFP
cGFP-CAT
Mt-GFP
Mt-GFP-CAT
Epoxo
MG132
GFP
kDa
250
150
100
75
SDS-resistant
aggregates
GFP
25
20
Actin
37
B
cGFP
cGFP-CAT
epoxo
Tom20
GFP
Merge
C
Mt-GFP
Mt-GFP-CAT

Figure S5

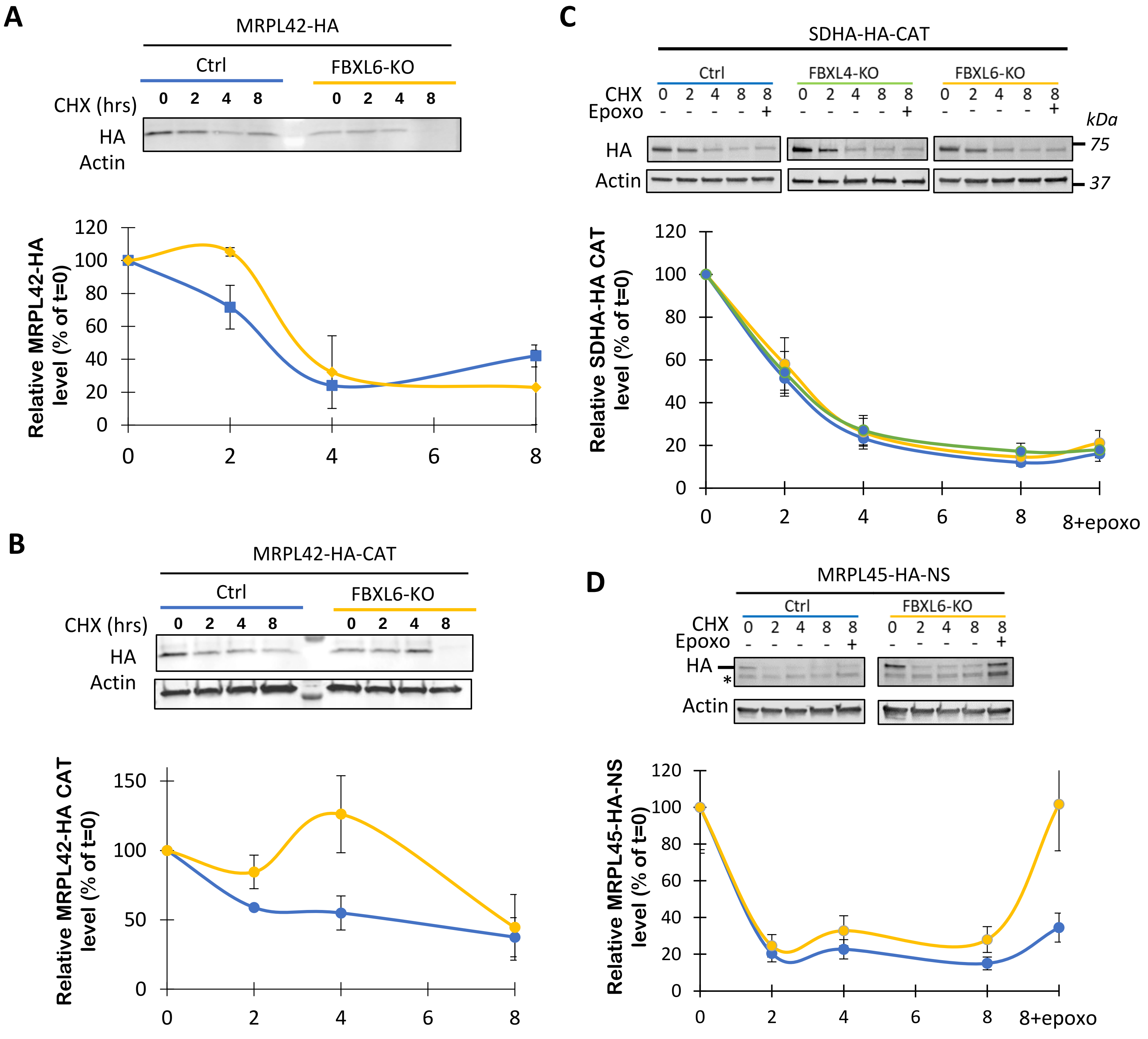
A
MRPL42-HA
Ctrl
FBXL6-KO
CHX (hrs)
0 2 4 8
0 2 4 8
HA
Actin
Relative MRPL42-HA level (% of t=0)
120
100
80
60
40
20
0
0 2 4 6 8
B
MRPL42-HA-CAT
Ctrl
FBXL6-KO
CHX (hrs)
0 2 4 8
0 2 4 8
HA
Actin
Relative MRPL42-HA CAT level (% of t=0)
150
100
50
0
0 2 4 6 8
C
SDHA-HA-CAT
Ctrl
FBXL4-KO
FBXL6-KO
CHX 0 2 4 8 8
Epoxo - - - - +
kDa
HA
Actin
75
37
Relative SDHA-HA CAT level (% of t=0)
120
100
80
60
40
20
0
0 2 4 6 8 8+epoxo
D
MRPL45-HA-NS
Ctrl
FBXL6-KO
CHX 0 2 4 8 8
Epoxo - - - - +
HA
*
Actin
Relative MRPL45-HA-NS level (% of t=0)
120
100
80
60
40
20
0
0 2 4 6 8 8+epoxo

### Figure S6

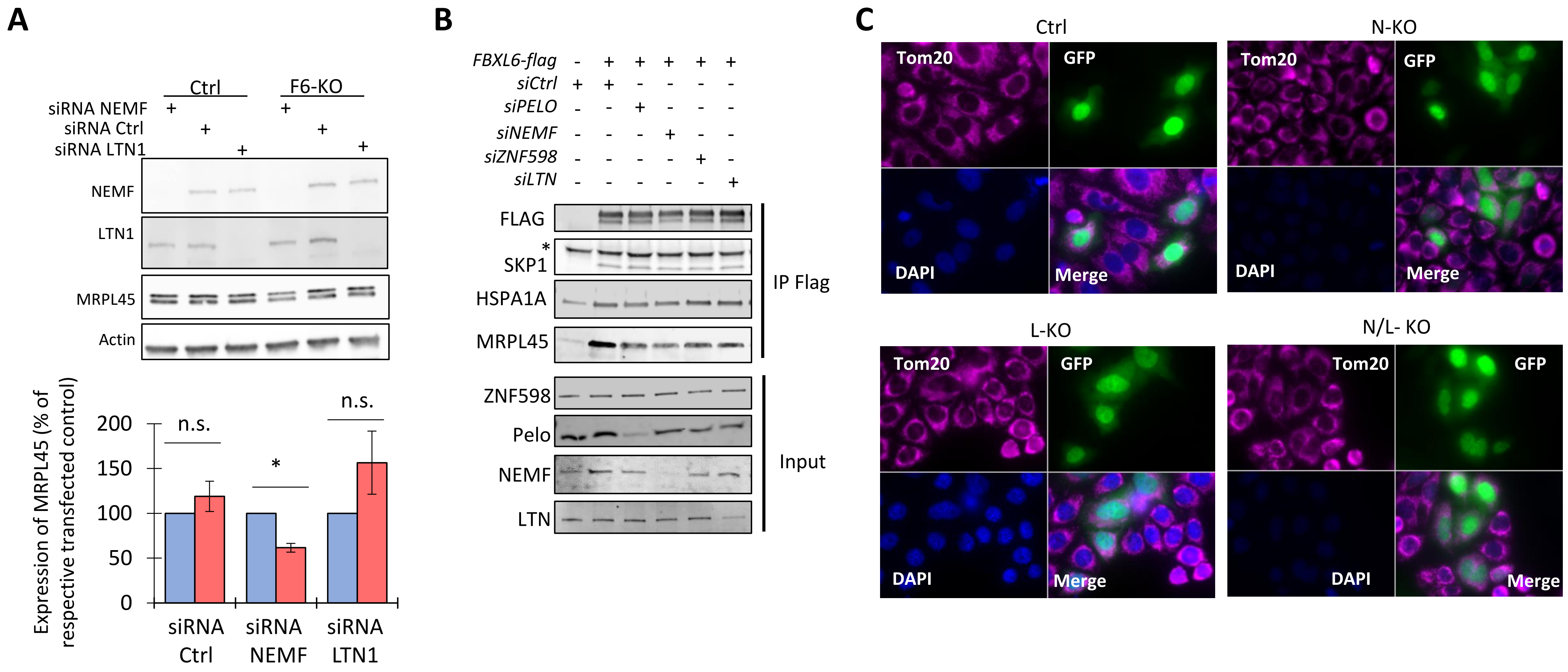

A
Ctrl
F6-KO
siRNA NEMF
siRNA Ctrl
siRNA LTN1
NEMF
LTN1
MRPL45
Actin
Expression of MRPL45 (% of respective transfected control)
200
150
100
50
0
n.s.
*
n.s.
siRNA Ctrl
siRNA NEMF
siRNA LTN1
B
FBXL6-flag
siCtrl
siPELO
siNEMF
siZNF598
siLTN
FLAG
SKP1
HSPA1A
MRPL45
IP Flag
ZNF598
Pelo
NEMF
LTN
Input
C
Ctrl
N-KO
L-KO
N/L- KO
Tom20
GFP
DAPI
Merge
